# Differential encoding of mammalian proprioception by voltage-gated sodium channels

**DOI:** 10.1101/2024.08.27.609982

**Authors:** Cyrrus M. Espino, Chetan Nagaraja, Serena Ortiz, Jacquelyn R. Dayton, Akash R. Murali, Yanki Ma, Emari L. Mann, Snigdha Garlapalli, Ross P. Wohlgemuth, Sarah E. Brashear, Lucas R. Smith, Katherine A. Wilkinson, Theanne N. Griffith

## Abstract

Animals that require purposeful movement for survival are endowed with mechanosensory neurons called proprioceptors that provide essential sensory feedback from muscles and joints to spinal cord circuits, which modulates motor output. Despite the essential nature of proprioceptive signaling in daily life, the mechanisms governing proprioceptor activity are poorly understood. Here, we have identified distinct and nonredundant roles for two voltage-gated sodium channels (NaVs), NaV1.1 and NaV1.6, in mammalian proprioception. Deletion of NaV1.6 in somatosensory neurons (NaV1.6^cKO^ mice) causes severe motor deficits accompanied by complete loss of proprioceptive transmission, which contrasts with our previous findings using similar mouse models to target NaV1.1 (NaV1.1^cKO^). In NaV1.6^cKO^ animals, loss of proprioceptive feedback caused non-cell- autonomous impairments in proprioceptor end-organs and skeletal muscle that were absent in NaV1.1^cKO^ mice. We attribute the differential contribution of NaV1.1 and NaV1.6 in proprioceptor function to distinct cellular localization patterns. Collectively, these data provide the first evidence that NaV subtypes uniquely shape neurotransmission within a somatosensory modality.

**Teaser:** Voltage gated sodium channels differentially encode mammalian proprioception via distinct cellular localization patterns.

## Introduction

Proprioception, often referred to as our “sixth sense”, is a largely unconscious sensation that allows for the detection of one’s own body position and movement in space (*1*, *2*). Proprioceptive signaling is initiated by a subclass of peripheral mechanosensory neurons, called proprioceptors, whose cell bodies reside in the dorsal root ganglia (DRG) or mesencephalic trigeminal nucleus (*1*, *3*, *4*). The peripheral axons of proprioceptors innervate skeletal muscle and form mechanosensitive end organs, referred to as muscle spindles and Golgi tendon organs, which are activated by changes is muscle length or force, respectively (*1*). In proprioceptors, the mechanosensitive ion channel Piezo2 transduces changes in muscle movement into electrical signals that give rise to sustained trains of action potentials, which are subsequently transmitted to spinal cord circuits (*4*, *5*). Indeed, patients harboring Piezo2 loss-of-function mutations have impaired proprioception in the absence of visual input (*6*). Recently, we determined that the voltage-gated sodium channel (NaV), NaV1.1, is also essential for mammalian proprioception, and plays a specific role in maintaining proprioceptor firing during sustained muscle stretch (*7*). Furthermore, we determined NaV1.1 to be haploinsufficient for proprioceptor function and motor behaviors, which is consistent with the clinical manifestations associated with the thousands of human disease-causing mutations associated with its gene, *Scn1a*. Surprisingly, NaV1.1 was not required for muscle proprioceptor responses to dynamic muscle movement or vibration. This raises the question as to whether NaVs play distinct roles in encoding proprioceptive signals.

In addition to NaV1.1, proprioceptors also express NaV1.6 and NaV1.7 (*7*). NaV1.7 is most notable for its role in pain signaling, whereby gain- or loss-of-function mutations in *Scn9a*, the gene encoding NaV1.7, cause congenital hypersensitivity or insensitivity to pain, respectively (*8*, *9*) . Mice and humans lacking NaV1.7, however, do not have reported motor deficits, indicating a limited role for this channel in proprioception at the behavioral level (*9*, *10*). Conversely, the gene encoding NaV1.6, *Scn8a*, is linked to various pathophysiological conditions associated with motor impairments, such as developmental epileptic encephalopathy and ataxia (*11*). Furthermore, global inactivation of *Scn8a* in mice leads to hind limb paralysis and death by postnatal day (P) 21 (*12*). In cerebellar Purkinje neurons, loss of NaV1.6 significantly reduces spontaneous activity and leads to impairments in motor coordination (*13*). While these data highlight critical roles for NaV1.6 in brain-mediated motor control, NaV1.6 function remains understudied in the peripheral nervous system, and how this channel contributes to proprioception is unknown. Importantly, understanding the unique contributions of NaVs to peripheral proprioception will enhance our mechanistic understanding of the sensorimotor phenotypes associated with various NaV channelopathies.

In the present study, we set out to determine whether NaVs plays distinct or redundant roles in proprioceptive signaling, focusing on the contributions of NaV1.1 and NaV1.6. The use of a *Pvalb- Cre* mouse line to drive NaV deletion in proprioceptors is not feasible due to parvalbumin expression in the brain and spinal cord (*7*, *14*, *15*). Thus, we used a somatosensory-neuron wide genetic targeting strategy to conditionally deleted NaV1.6 (*Pirt^Cre/+^;Scn8a^fl/fl^*, NaV1.6^cKO^) and found this resulted in severe impairments in motor coordination that were phenotypically distinct from those we previously observed in mice lacking NaV1.1 in somatosensory neurons (*Pirt^Cre/+^;Scn1a^fl/fl^*, NaV1.1^cKO^, *7*). In line with behavioral observations, *ex vivo* proprioceptor muscle-nerve recordings showed neurotransmission in response to both dynamic and static muscle movement was abolished in the absence of NaV1.6, which contrasts with our prior finding of a selective role for NaV1.1 in proprioceptor encoding of static muscle stretch. Electrophysiological recordings of the proprioceptor-mediated monosynaptic reflex in the spinal cord further confirmed an essential, albeit developmentally dependent, role for NaV1.6 in proprioceptor synaptic function, whereas NaV1.1 was found to be dispensable. NaV1.6^cKO^ mice also exhibited abnormal muscle spindle end organ structure, which was not observed in NaV1.1^cKO^ mice, suggesting severely but not moderately impaired proprioceptive signaling interferes with proprioceptor end organ development. Surprisingly, we also observed non-cell-autonomous deficits in skeletal muscle development in NaV1.6^cKO^ mice, but not NaV1.1^cKO^ mice, that are suggestive of blocked hypertrophy during development. Finally, cellular localization experiments found NaV1.1 and NaV1.6 occupy discrete excitable domains in proprioceptor muscle spindles, which we predict underlies their unique roles in electrical transmission.

Collectively, our findings reveal that NaV1.1 and NaV1.6 are both essential to proprioceptive signaling but have independent and non-redundant functions. The differential contribution of NaV1.1 and NaV1.6 to the activity of individual somatosensory neuron subtypes has not been investigated, and we hypothesize our findings are broadly applicable to other somatosensory neurons that rely on these channels for neuronal signaling.

## Results

### Genetic ablation of NaV1.6 in sensory neurons leads to profound motor coordination deficits

To examine the *in vivo* role of NaV1.6 in sensory-driven motor behaviors, we generated a mouse line in which NaV1.6 is deleted in all peripheral sensory neurons: *Pirt^Cre/+^Scn8a^fl/fl^*(hereafter referred to as NaV1.6^cKO^), an approach we previously used to investigate NaV1.1 function in proprioception (*7*). While not selective to proprioceptors, this approach avoids significant off-target effects on the central nervous system, which include premature death and seizures (*14*). NaV1.6^cKO^ mice displayed extreme motor deficits that were absent in mice retaining a single copy (NaV1.6^het^) or both copies (NaV1.6^fl/fl^) of *Scn8a* (Fig. 1). Motor deficits included abnormal hind limb position when suspended by the tail (Fig. 1 A to C, top; movie S1) or when placed on a flat surface (Fig. 1 A to C, bottom; movie S2) and an inability to use the tail to guide movements. The motor phenotype produced by NaV1.6 deletion was more severe than the phenotype we observed following deletion of NaV1.1 in sensory neurons (*Pirt^Cre/+^;Scn1a^fl/fl^,* NaV1.1^cKO^, *7*). Interestingly, however, NaV1.6^cKO^ mice did not display the tremor-like movements we previously observed in NaV1.1^cKO^ animals, highlighting a behaviorally distinct phenotype between the two models. We quantified spontaneous locomotion in the open-field and found that NaV1.6^cKO^ animals traveled significantly less distance (Fig. 1D) and were slower (Fig. 1E) compared to NaV1.6^het^ and NaV1.6^fl/fl^ animals. There were no genotype dependent differences in time spent moving (Fig. 1F), suggesting that motivation to move is not impaired in NaV1.6^cKO^ mice. Time spent in the center was also not different between genotypes (Fig. S1, A). Furthermore, we did not observe any sex-dependent differences between genotypes (Fig. S1 B to D). Using a grip strength meter, we quantified grip force when all four paws were placed on a metal grid and found that NaV1.6^cKO^ animals had a significantly reduced grip strength compared to other genotypes (Fig. 1G). We next assessed motor coordination using the rotarod; however, the severe motor phenotype of NaV1.6^cKO^ mice precluded their testing in this assay. We did not observe genotype dependent differences in latency to fall between NaV1.6^het^ and NaV1.6^fl/fl^ animals across training days (Fig. 1H) or on the final day of testing (Fig. 1I). Collectively, these data show that genetic ablation of NaV1.6 in sensory neurons leads to severe motor deficits that are distinct to those due to NaV1.1 deletion. Interestingly, we previously reported that NaV1.1 was haploinsufficient in sensory neurons for motor behaviors; however, these results suggest a single copy of NaV1.6 is sufficient to drive normal motor function at the behavioral level. Nevertheless, we did observe NaV1.6 haploinsufficiency at the afferent level.

**Fig. 1.**
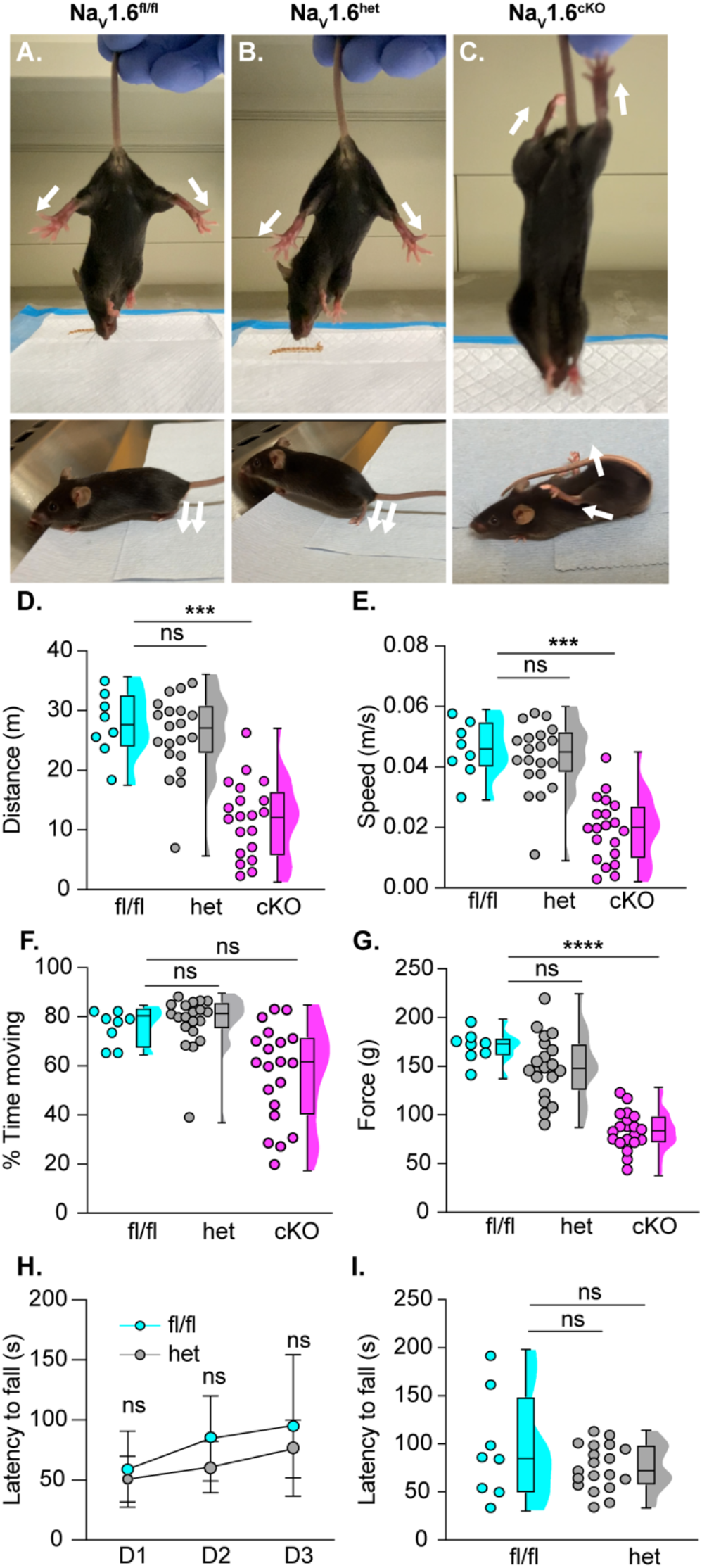
NaV1.6 is required for somatosensory neuron-driven motor behaviors and function. Representative images showing limb position of adult NaV1.6^fl/fl^ (**A**), NaV1.6^het^ (**B**), and NaV1.6^cKO^ (**C**) mice suspended from the tail (above) and on flat surface (below). White arrows indicate the direction of hind limbs. Quantification of distance traveled (**D,** NaV1.6^het^ p > 0.999, NaV1.6^cKO^ p = 0.0003, compared to NaV1.6^fl/fl^), average speed (**E,** NaV1.6^het^ p > 0.999, NaV1.6^cKO^ p = 0.0003), and the percent of time spent moving (**F,** NaV1.6^het^ p > 0.999, NaV1.6^cKO^ p = 0.05, compared to NaV1.6^fl/fl^) for NaV1.6^fl/fl^ (cyan), NaV1.6^het^ (grey), and NaV1.6^cKO^ (magenta) mice as measured in the open field assay for a 10-minute testing period. (**G**) Average grip force in grams measured across 6 consecutive trials; NaV1.6^het^ p = 0.4947, NaV1.6^cKO^ p < 0.0001, compared to NaV1.6^fl/fl^. (**H**) Average latency to fall from the rotarod across three consecutive training days. No statistically significant genotype-dependent difference was observed (p = 0.1342). (**I**) Average latency to fall on third day of testing (NaV1.6^het^ p = 0.3943, compared to NaV1.6^fl/fl^). Each dot represents one animal, except in (**H**) were each dot represents the mean across animals. Box and whisker plots represent maximum, minimum, median, upper and lower quartiles of data sets. A Kruskal-Wallis test with Dunn’s multiple comparisons (**D** to **G**), a Two-way ANOVA with Sidak’s multiple comparisons (**H**), and a Welch’s T-test (**I**) were used to determine statistical significance. NaV1.6^fl/fl^ N = 8, NaV1.6^het^ N=20, NaV1.6^cKO^ N=20.

### NaV1.6 is required for transmission of proprioceptive signals from muscle spindle afferents

Our prior work found that NaV1.1 plays a key role maintaining muscle afferent activity only during static muscle stretch (*7*). Deletion of NaV1.1 had no effect on muscle afferent responses to dynamic muscle movement or vibratory stimuli. To test whether NaV1.6 serves a similarly specific role in proprioceptive transmission, we used an *ex vivo* muscle nerve preparation to investigate proprioceptor activity from muscle spindle afferents (*16*). First, we tested afferent firing in response to a series of ramp and hold stretches (Fig. 2). Afferents from NaV1.6^fl/fl^ mice displayed consistent firing throughout the duration of a 4 s ramp and hold stretch protocol and had a high likelihood of resting discharge (Fig. 2A), consistent with wild-type Group Ia and II proprioceptor responses. Afferents from NaV1.6^het^ mice had a similar prevalence of resting discharge compared to NaV1.6^fl/fl^ mice and did not exhibit any significant differences in firing during ramp and hold stretches (Fig. 2B). Strikingly, afferents from NaV1.6^cKO^ mice never possessed resting discharge and neurotransmission during ramp and hold stretches were nearly abolished (Fig. 2C). In 6 out of the 10 mice tested, no stretch-sensitive electrical activity was observed despite the muscle exhibiting healthy twitch contractions. We quantified afferent properties by examining instantaneous firing frequencies and found a significant reduction in firing in NaV1.6^cKO^ afferents compared to NaV1.6^fl/fl^ afferents across all stretch lengths (Fig. 2D). There were no significant genotype dependent differences in firing between NaV1.6^het^ and NaV1.6^fl/fl^ afferents. We also quantified the regularity of afferent firing by measuring the coefficient of variation of the interspike interval (ISI CV). ISI CV was similar between NaV1.6^fl/fl^ and NaV1.6^het^ afferents but was significantly higher in NaV1.6^cKO^ afferents (Fig. 2E). Together these findings provide strong evidence that NaV1.6 is required for proprioceptor encoding of static stretch.

**Fig. 2.**
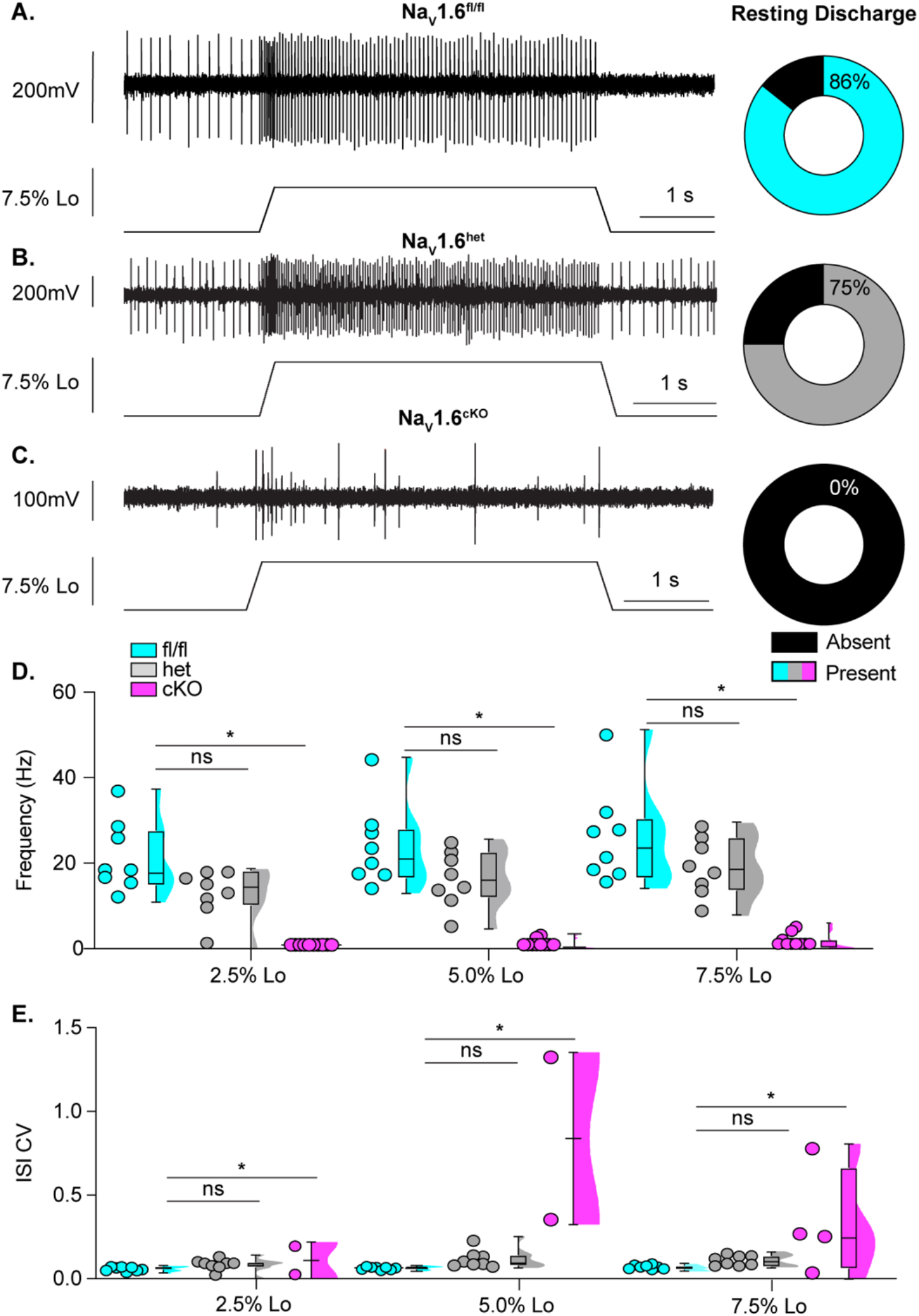
Loss of NaV1.6 abolishes muscle proprioceptor static stretch sensitivity. Representative responses to ramp-and-hold muscle stretch at 7.5% of optimal length (Lo) in NaV1.6^fl/fl^ (**A**), NaV1.6^het^ (**B**), and NaV1.6^cKO^ (**C**) muscle proprioceptors. The percentage of afferents that displayed resting discharge at Lo are represented by the pie charts to the right (black indicates absence of resting discharge). (**D**) Quantification of afferent firing frequency 3.25 to 3.75 seconds into stretch protocol. NaV1.6^fl/fl^ (cyan), NaV1.6^het^ (grey), and NaV1.6^cKO^ (magenta). NaV1.6^het^ p = 0.178, NaV1.6^cKO^ p = 0.001, compared to NaV1.6^fl/fl^. (**E**) Firing regularity was quantified as the coefficient of variation of the interspike interval (ISI CV) 1.5 to 3.5 seconds into the stretch protocol. NaV1.6^het^ p = 0.669, NaV1.6^cKO^ p = 0.000, compared to NaV1.6^fl/fl^. In 6 out of 10 animals we observed no response to stretch and therefore could only include the quantifiable responses from 4 afferents from NaV1.6^cKO^ mice. Only quantifiable responses were included in statistical analyses in **D** and **E**. Box and whisker plots represent maximum, minimum, median, upper and lower quartiles of data sets. Each dot represents a single afferent. A two-way mixed-design ANOVA (Dunnett’s post-hoc comparison) was used to determine statistical significance in **D** and **E**. NaV1.6^fl/fl^ n = 8, N=7; NaV1.6^het^ n=8, N=8; NaV1.6^cKO^ n=4, N=10. n = afferents, N = mice.

Given that NaV1.1 only contributes to proprioceptor afferent firing in response to static muscle stretch, we next asked whether this was also true for NaV1.6. Afferents were tested using a series of sinusoidal vibration protocols at varying frequencies and stimulus amplitudes. In line with the absence of electrical activity during static stretch, NaV1.6 afferents were completely unable to entrain to vibratory stimuli regardless of stimulus amplitude and frequency (Fig. 3). Interestingly, logistic regression analyses of entrainment probability found that compared to NaV1.6^fl/fl^ afferents, afferents from NaV1.6^het^ animals were significantly less likely to entrain sinusoidal waves (p<0.001; Fig. 3 A and B). This analysis could not be used to assess entrainment probability in NaV1.6^cKO^ afferents because these afferents never entrained to vibration. Consistent with logistic regression analyses, quantification of the instantaneous firing frequency at 25μm amplitude vibrations found significant impairments in the ability of NaV1.6^het^ afferents to respond to vibration, consistent with the notion that NaV1.6 is partially haploinsufficient at the proprioceptor afferent level (Fig. 3D). Thus, unlike NaV1.1 which only serves a role in maintaining proprioceptor responses to static stretch, we find NaV1.6 plays a direct role in transmitting both dynamic and static muscle movement.

**Fig 3.**
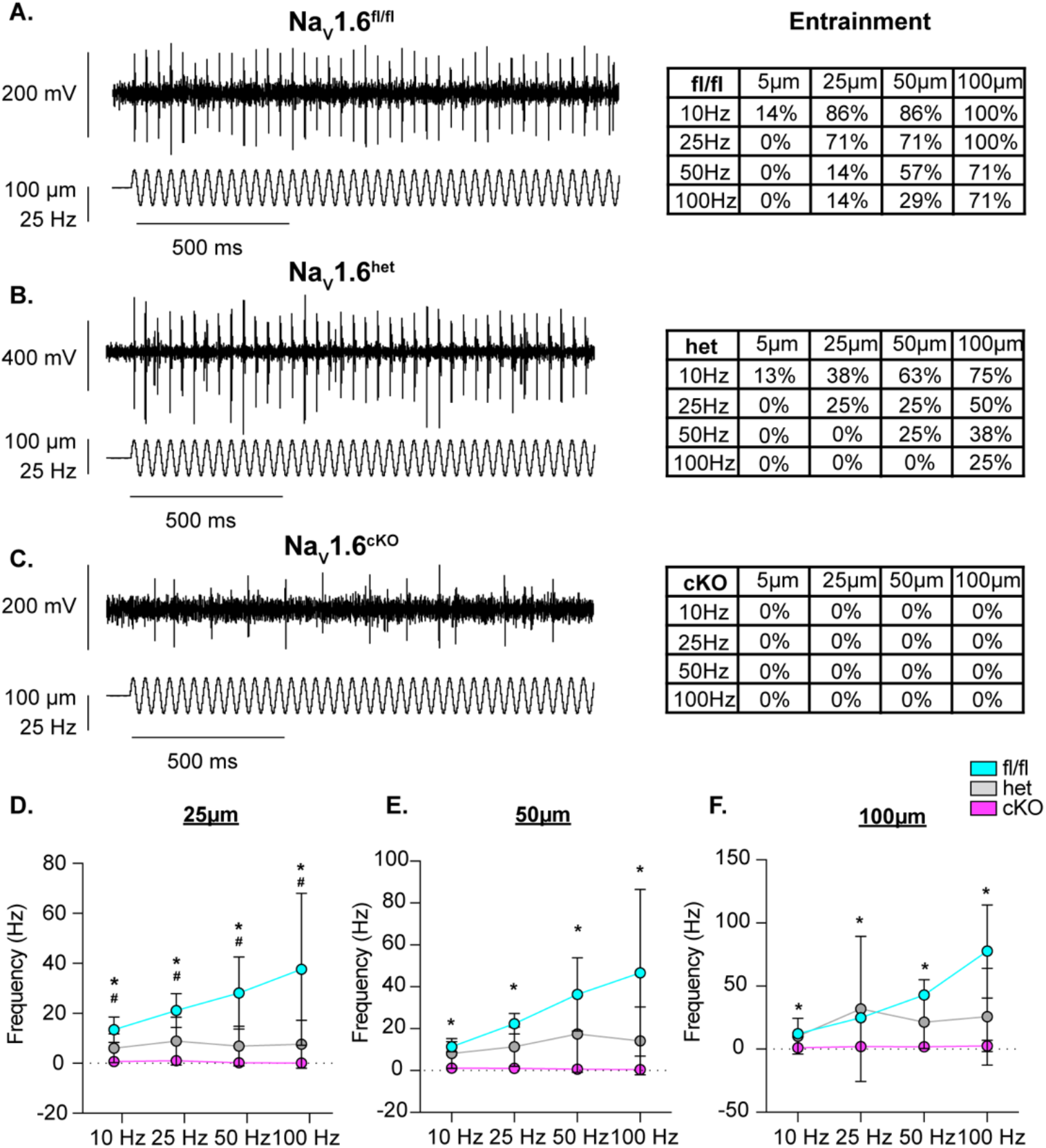
NaV1.6 is required for proprioceptor responses to vibration. Representative traces from NaV1.6^fl/fl^ (**A**), NaV1.6^het^ (**B**), and NaV1.6^cKO^ (**C**) afferents that were able to entrain to a 25 Hz, 100 μm amplitude vibration stimulus. Tables to the right indicate the percentage of afferents that were able to entrain across stimulus frequencies and amplitudes (NaV1.6^fl/fl^, top; NaV1.6^het^, middle; NaV1.6^cKO^, bottom; **D** to **E**) Quantification of firing frequency across vibration amplitudes. At 25μm (**D**) NaV1.6^het^ p = 0.005 (# denotes significance in NaV1.6^het^), NaV1.6^cKO^ p = 0.001, compared to NaV1.6^fl/fl^ (* denotes significance in NaV1.6^cKO^). At 50μm (**E**) NaV1.6^het^ p = 0.053, NaV1.6^cKO^ p = 0.002, compared to NaV1.6^fl/fl^. At 100μm (**F**) NaV1.6^het^ p = 0.414, NaV1.6^cKO^ p = 0.018 compared to NaV1.6^fl/fl^. NaV1.6^fl/fl^ (cyan), NaV1.6^het^ (grey), and NaV1.6^cKO^ (magenta). A two-way mixed- design ANOVA (Dunnett’s post-hoc comparison) was used to determine statistical in **D** to **F**. Box and whisker plots represent maximum, minimum, median, upper and lower quartiles of data sets. Each dot represents the average afferent response per genotype. NaV1.6^fl/fl^ n = 8, N=7; NaV1.6^het^ n=8, N=8; and NaV1.6^cKO^ n= 4, N=10. n=afferents, N=mice.

### NaV1.6 is essential for proprioceptor synaptic transmission in a developmentally dependent manner

Electrical signals initiated at proprioceptive end organs in skeletal muscle are transmitted to central circuits in the spinal cord. Specifically, proprioceptor Ia afferents directly synapse with alpha motor neurons, comprising the monosynaptic reflex response (*17*, *18*). This spinal circuit provides a tractable model to assess proprioceptor synaptic transmission. Our current results demonstrate that NaV1.6 plays a central role in sensory transmission from muscle spindles; thus, we next asked whether the peripheral deficits we observed in *ex vivo* muscle nerve recordings are also evident in proprioceptive circuits in the spinal cord. We used an *ex vivo* hemisected spinal cord preparation and measured properties of the monosynaptic reflex circuit in NaV1.6^fl/fl^, NaV1.6^het^, and NaV1.6^cKO^mice (Fig. 4). We first analyzed responses from mice in early postnatal development (P6 to P11) as all prior work has used this age range for monosynaptic reflex analysis, largely due to technical challenges associated with increased myelination in the ventral horn as development proceeds (*19*). In stark contrast to our muscle-nerve recordings, monosynaptic responses were similar between genotypes at this timepoint (Fig. 4). We only observed a significant difference in response latency in NaV1.6^cKO^ hemicords compared to NaV1.6^het^ and NaV1.6^fl/fl^ hemicords (Fig. 4B). No other genotype dependent differences were observed across quantified parameters (Fig. 4 C to F). These findings suggest that during early postnatal development, NaV1.6 is dispensable for proprioceptor synaptic transmission.

**Fig. 4.**
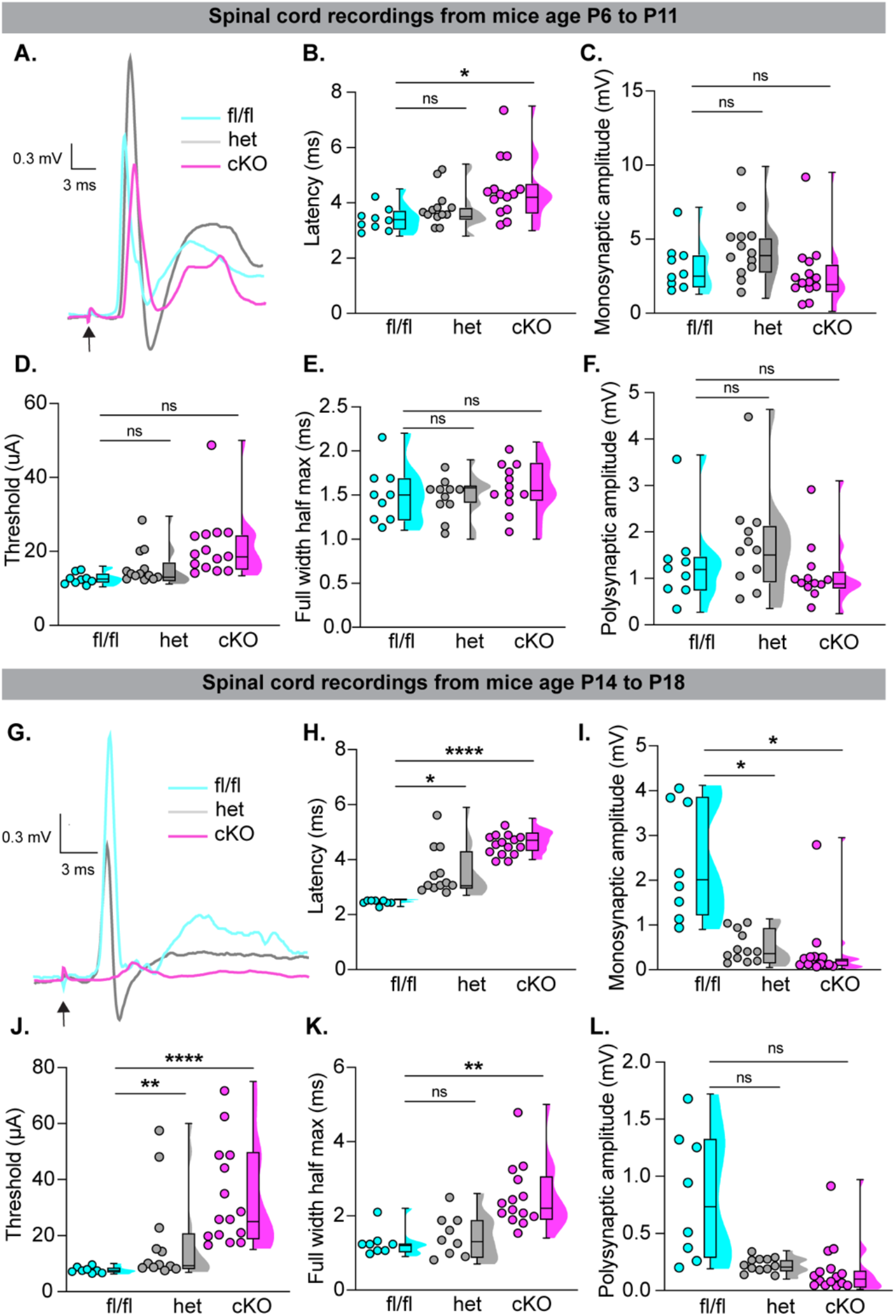
NaV1.6 plays a developmentally dependent role in proprioceptor synaptic transmission in the spinal cord. (**A**) Representative monosynaptic reflex responses from NaV1.6^fl/fl^ (cyan), NaV1.6^het^ (grey), and NaV1.6^cKO^ (magenta) hemicords during postnatal days 6 to 11. Quantification of response properties. (**B**) Response latency, NaV1.6^het^ p = 0.760, NaV1.6^cKO^ p = 0.019, compared to NaV1.6^fl/fl^. (**C**) Monosynaptic response amplitude, NaV1.6^het^ p = 0.238, NaV1.6^cKO^ p = 0.640, compared to NaV1.6^fl/fl^. (**D**) Stimulus threshold, NaV1.6^het^ p = 0.910, NaV1.6^cKO^ p = 0.271, compared to NaV1.6^fl/fl^. (**E**) Full width half max, NaV1.6^het^ p = 0.999, NaV1.6^cKO^ p = 0.929, compared to NaV1.6^fl/fl^. (**F**) Polysynaptic response amplitude, NaV1.6^het^ p = 0.514, NaV1.6^cKO^ p = 0.704, compared to NaV1.6^fl/fl^. (**G**) Representative monosynaptic reflex responses in NaV1.6^fl/fl^ (cyan), NaV1.6^het^ (grey), and NaV1.6^cKO^ (magenta) hemicords during postnatal days 14 to 18. (**H**) Response latency, NaV1.6^het^ p = 0.023, NaV1.6^cKO^ p < 0.0001, compared to NaV1.6^fl/fl^. (**I**) Monosynaptic response amplitude, NaV1.6^het^ p = 0.037, NaV1.6^cKO^ p = 0.018, compared to NaV1.6^fl/fl^. (**J**) Stimulus threshold, NaV1.6^het^ p = 0.164, NaV1.6^cKO^ p <0.0001, compared to NaV1.6^fl/fl^. (**K**) Full width half max, NaV1.6^het^ p = 0.784, NaV1.6^cKO^ p <0.0001, compared to NaV1.6^fl/fl^. (**L**) Polysynaptic response amplitude, NaV1.6^het^ p = 0.143, NaV1.6^cKO^ p = 0.092, compared to NaV1.6^fl/fl^. Each dot represents a single hemicord. (**A** to **F**) NaV1.6^fl/fl^ n=9, NaV1.6^het^ n=13, and NaV1.6^cKO^ n=14. (**G** to **L**) NaV1.6^fl/fl^ n=8, NaV1.6^het^ n=12, and NaV1.6^cKO^ n=15. N=8-15. n=hemicords, N=mice. Box and whisker plots represent maximum, minimum, median, upper and lower quartiles of data sets. A two-way mixed- design ANOVA (Tukey’s post-hoc comparison) was used to determine statistical significance.

Interestingly, previous studies indicate that proprioceptors are not transcriptionally mature until walking behaviors begin to emerge (*20*), which occurs around P13. Thus, we decided to test monosynaptic responses beyond this time point (P14 to P18). To our knowledge, this is first systematic analysis of the monosynaptic reflex this late in postnatal development. Strikingly, at this age, proprioceptive synaptic transmission is nearly lost in NaV1.6^cKO^ hemicords (Fig. 4G). We found highly significant genotype dependent differences between NaV1.6^cKO^ and NaV1.6^fl/fl^ hemicords across all quantified parameters (Fig. 4 H to L). This suggests that following the onset of walking behaviors, NaV1.6 is essential for proprioceptor synaptic transmission onto alpha motor neurons, which is also consistent data from *ex vivo* muscle nerve recordings in adult afferents (Figs. 2 and 3). Additionally, we also found significantly increased monosynaptic reflex response latencies and thresholds (Fig. 4H and J), as well as significantly reduced response amplitudes in NaV1.6^het^ hemicords compared to NaV1.6^fl/fl^ controls (Fig. 4 I). Finally, when looking within genotypes, we found that unlike in NaV1.6^fl/fl^ hemicords, which only showed a significant increase in response latency, there was a significant degradation of the monosynaptic reflex response in NaV1.6^het^ and NaV1.6^cKO^ hemicords (Table S1). Thus, during postnatal development, NaV1.6^fl/fl^ animals exhibit an enhancement in central proprioceptive signaling, whereas in both NaV1.6^het^ and NaV1.6^cKO^ mice, central proprioceptive signaling degrades. This provides additional evidence that at the circuit level, NaV1.6 is haploinsufficient for proprioceptor synaptic function.

The developmental dependence of the proprioceptor mediated monosynaptic reflex response on NaV1.6 prompted us to investigate motor behaviors in this line at P7 and P14, before and after the onset of weight bearing locomotion, respectively (Fig. S2). We analyzed P7 mice in a righting reflex assay, a hindlimb suspension test, a grasping reflex assay, and quantified hindlimb angle. In line with spinal cord electrophysiology data, behavioral testing at P7 found a minimal role of NaV1.6 across behavioral assays (Fig. S2, A to H); we only observed a significant difference in hindlimb angle at this age. Conversely, when we analyzed motor abilities in a limb coordination assay at P14, we observed significant differences in functional grasping in both NaV1.6^het^ and NaV1.6^cKO^ mice compared to NaV1.6^fl/fl^ controls (Fig. S2, I to J). These data highlight a developmentally-specific contribution of NaV1.6 to proprioceptive synaptic transmission in the spinal cord, which also manifests at the behavioral level.

Because we did not observe a role for NaV1.6 in the proprioceptor mediated monosynaptic reflex response between P6 and P11, we next asked whether instead NaV1.1 was required for proprioceptor synaptic transmission at this developmental stage. In line with our findings across NaV1.6 genotypes, monosynaptic reflex responses were genotype-independent in the NaV1.1 mouse line in this age range (Fig. 5). These data suggest that proprioceptor synaptic transmission at early postnatal development is not dependent on NaV1.1 or NaV1.6 and could indicate NaV functional redundancy in proprioceptors prior to the onset of walking behaviors. We attempted to measure the monosynaptic reflex in later postnatal development (P14 to P18) in these mice; however, responses across all genotypes were too small to reliably quantify, despite our ability to obtain recordings at this age from mice in the Nav1.6-line (Fig. 4) as well as age matched C57Bl/6J controls (Fig. S3). Thus, we cannot rule out the possibility that NaV1.1 may serve a role in proprioceptor mediated synaptic transmission following the acquisition of weight bearing locomotion. Behavioral testing at P7 did not reveal changes in motor function across genotypes (Fig. S4 A to B). At P14 however, we did observe a significant difference in functional grasping in NaV1.1^cKO^ mice compared to NaV1.1^fl/fl^ controls (Fig. S4E), suggesting NaV1.1 is also becomes required for motor coordination at the onset of walking behaviors. These data show that in early postnatal development neither NaV1.6 or NaV1.1 alone is required for proprioceptor synaptic transmission; however, upon the acquisition of weight-bearing locomotion, we find that NaV1.6 becomes functionally dominant at the circuit level, and both NaV1.1 and NaV1.6 are required at the behavioral level.

**Fig. 5.**
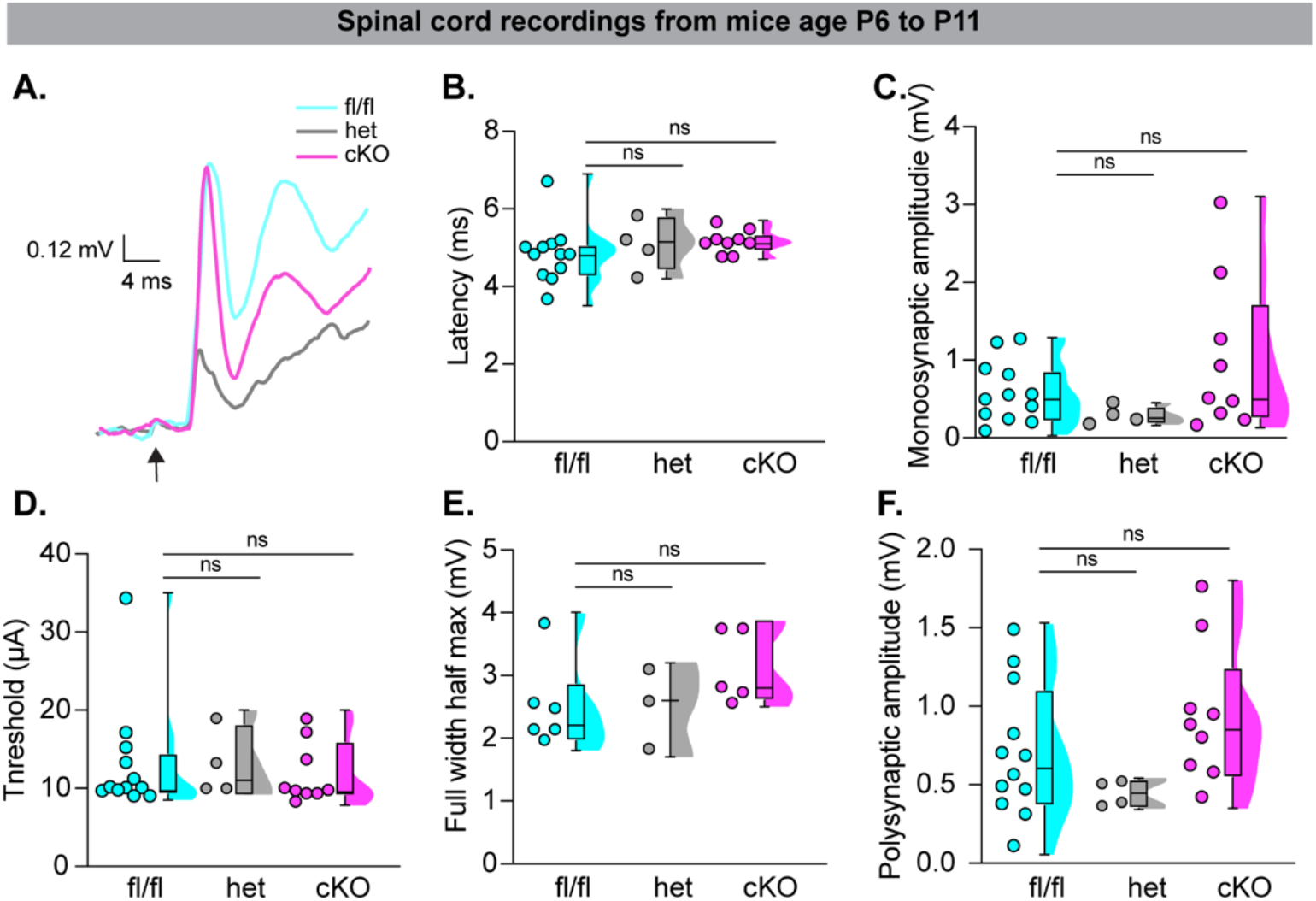
NaV1.1 does not contribute to proprioceptor synaptic transmission prior to the onset of walking behaviors. (**A**) Representative monosynaptic reflex responses from NaV1.1^fl/fl^ (cyan), NaV1.1^het^ (grey), and NaV1.1^cKO^ (magenta) hemicords recorded during postnatal days 6 to 11. (**B** to **D**) Quantification of monosynaptic response properties. (**B**) Response latency, NaV1.1^het^ p = 0.723, NaV1.1^cKO^ p = 0.238, compared to NaV1.1^fl/fl^. (**C**) Monosynaptic response amplitude, NaV1.1^het^ p = 0.378, NaV1.1^cKO^ p > 0.999, compared to NaV1.1^fl/fl^. (**D**) Stimulus threshold, NaV1.1^het^ p>0.999, NaV1.1^cKO^ p>0.999, compared to NaV1.1^fl/fl^. (**E**) Full width half max, NaV1.1^het^ p>0.999, NaV1.1^cKO^ p = 0.255, compared to NaV1.1^fl/fl^. (**F**) Polysynaptic response amplitude, NaV1.1^het^ p = 0.574, NaV1.1^cKO^ p = 0.473, compared to NaV1.1^fl/fl^. Each dot represents a single hemicord. NaV1.1^fl/fl^ n=12, NaV1.1^het^ n=4, and NaV1.1^cKO^ n=9. N=4-10 mice. Box and whisker plots represent maximum, minimum, median, upper and lower quartiles of data sets. A Kruskal-Wallis test with Dunn’s multiple comparisons was used to determine statistical significance.

### Severely, but not moderately, impaired proprioception results in deficits in muscle spindle development

A recent study found that loss of Piezo2 or NaV1.6 in sensory neurons led to changes in tactile sensory neuron end organ development (*21*). This study raised the possibility that NaV1.6 may also regulate muscle spindle development. As with our prior examination of NaV1.1^cKO^ mice compared to controls, we did not observe a reduction in the overall number of proprioceptors in DRG sections (identified by *Pvalb* and *Runx3* colocalization) between NaV1.6^fl/fl^, NaV1.6^het^, and NaV1.6^cKO^ mice (Fig. S5 B). We next examined the structure of muscle spindles in NaV1.6^cKO^ mice by performing immunohistochemistry against vesicular glutamate transporter 1 (VGLUT1) to visualize muscle spindle sensory wrappings in sections of extensor digitorum longus muscle. Qualitative observation of muscle spindles from NaV1.6^cKO^ mice show striking structural abnormalities in sensory wrappings that were not present in NaV1.6^fl/fl^ or NaV1.6^het^ animals (Fig. 6 A to C). To validate that the structural changes we observed were not due disruptions in VGLUT1 expression, a subset of experiments were conducted with both VGLUT1 and the pan-neuronal marker βIII-tubulin. Both antibodies showed highly similar levels of immunoreactivity, indicating VGLUT1 is a good proxy for muscle spindle structure (Fig. S6). To our knowledge there is no standardized method to quantitatively assess the structure of muscle spindles. Thus, we devised a quantitative method to examine muscle spindle sensory terminals by measuring the colocalization of VGLUT1^+^ sensory wrappings around clusters of DAPI-positive nuclei, which represent intrafusal muscle fibers. By normalizing the number of wrappings to muscle spindle length, we calculated a wrapping efficiency index. We found that compared to NaV1.6^fl/fl^ and NaV1.6^het^ animals, muscle spindles from NaV1.6^cKO^ mice had significantly reduced wrapping efficiency indices, demonstrating that NaV1.6 in sensory neurons is required for muscle spindle development (Fig. 6D). In our prior work, we qualitatively reported that loss of NaV1.1 does not change muscle spindle structure (*7*). To confirm our previous findings using this quantitative approach, we analyzed the wrapping efficiency index of muscle spindles from NaV1.1^cKO^ mice compared to NaV1.1^het^ and NaV1.1^fl/fl^ controls (Fig. 6 E to H). In line with our previous work, NaV1.1 is not required for muscle spindle development as wrapping efficiency indices were not significantly different between genotypes (Fig. 6H). Thus, the above results show that muscle spindle development is impaired when proprioceptive signaling is severely, but not moderately, disrupted.

**Fig. 6.**
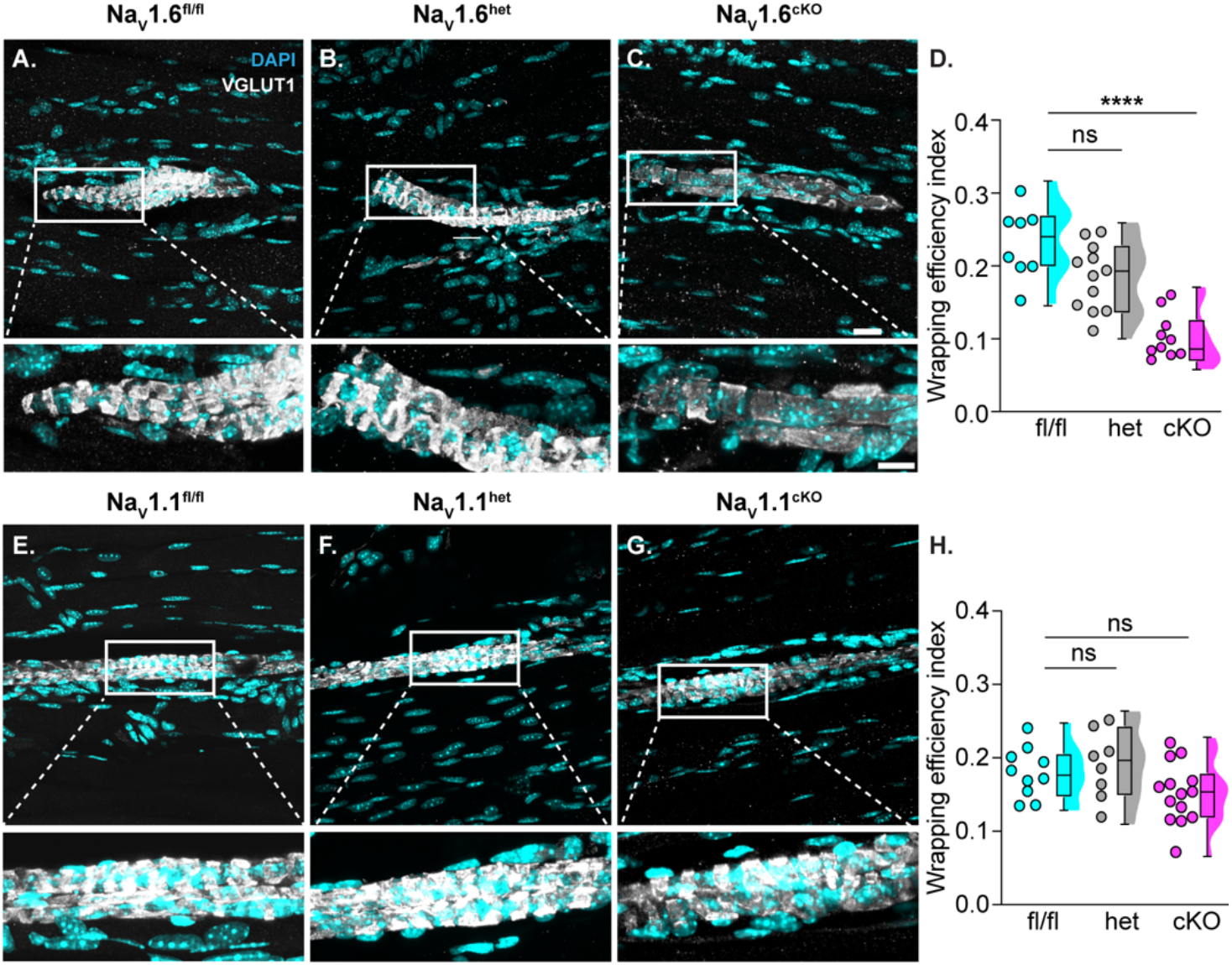
Electrical signaling deficits associated with NaV1.6- but not NaV1.1-deletion impairs muscle spindle development. Representative confocal images of muscle spindles from (**A**) NaV1.6^fl/fl^, (**B**) NaV1.6^het^, and (**C**) NaV1.6^cKO^ extensor digitorum longus (EDL) muscle sections (30 μm). Images were acquired with a 60x oil 1.4 NA lens. VGLUT1 (grey scale) labels proprioceptor sensory terminals and DAPI (cyan) labels nuclei. Insets below images show the colocalization of sensory terminals with DAPI. (**D**) Quantification of wrapping efficiency index based on colocalization of DAPI with VGLUT1. NaV1.6^het^ p = 0.0658, and NaV1.6^cKO^, p < 0.0001, compared to NaV1.6^fl/fl^. Representative images of muscle spindles from (**E**) NaV1.1^fl/fl^, (**F**) NaV1.1^het^, (**G**) NaV1.1^cKO^. (**H**) Quantification of wrapping efficiency index based on colocalization of DAPI with VGLUT1. NaV1.1^het^, p = 0.762, and NaV1.1^cKO^, p = 0.282, compared to NaV1.1^fl/fl^. Each dot represents a single muscle spindle section. (**D**) NaV1.6^fl/fl^ n=8, NaV1.6^het^ n=12, and NaV1.6^cKO^ n=10. (**H**) NaV1.1^fl/fl^ n=10, NaV1.1^het^ n=8, and NaV1.1^cKO^ n=14. N=3 mice per genotype. Box and whisker plots represent maximum, minimum, median, upper and lower quartiles of data sets. A one-way ANOVA (Dunnett’s post-hoc comparison) was used to determine statistical significance. Scale bar = 20 μm. Inset scale bar = 10 μm.

### Proprioceptive feedback is required for normal skeletal muscle development

Global inactivation of NaV1.6 leads to severe motor impairments accompanied by atrophy of skeletal muscle (*22*). It was hypothesized that these deficits were caused by loss of signal transmission from motor neurons; however, whether impaired proprioceptive feedback onto motor neurons is sufficient to impair skeletal muscle development has not been directly investigated. To address this, we analyzed skeletal muscle anatomy and function in the NaV1.6 mouse line. We collected soleus muscle from mice and labeled for slow (Type I) and fast (Type IIa) twitch muscle fibers. Muscle fibers from NaV1.6^cKO^ mice had a visible reduction in muscle fiber size compared to fibers from NaV1.6^het^ and NaV1.6^fl/fl^ mice (Fig. 7 A to C). To quantify these changes, we took an unbiased approach and measured muscle fiber properties using a semi-automatic muscle fiber analysis software in MATLAB (*23*). In agreement with qualitative observation, muscle fibers from NaV1.6^cKO^ mice displayed a significant decrease in fiber area compared to NaV1.6^het^ and NaV1.6^fl/fl^muscle (Fig. 7 D). A cumulative distribution plot shows the spread of muscle fiber area across genotypes (Fig. 7E). The proportion of Type I and Type IIa fibers were also similar between genotypes and were within the expected percentages for soleus muscle in wildtype animals (Fig. S7 A and B, *24*, *25*). Furthermore, we found that the size of both Type I and Type IIa fibers in NaV1.6^cKO^ mice were significantly reduced compared to other genotypes, indicating the changes in muscle fiber diameter and area were not fiber-type specific (Fig S7 C and D). We next examined whether the developmental changes in muscle fiber properties corresponded to alterations in intrinsic muscle strength and fatiguability; however, we found no differences in muscle function across genotypes (Fig. 7 F to I). Collectively, these reveal a non-cell autonomous role for proprioceptive feedback in skeletal muscle development.

**Fig 7.**
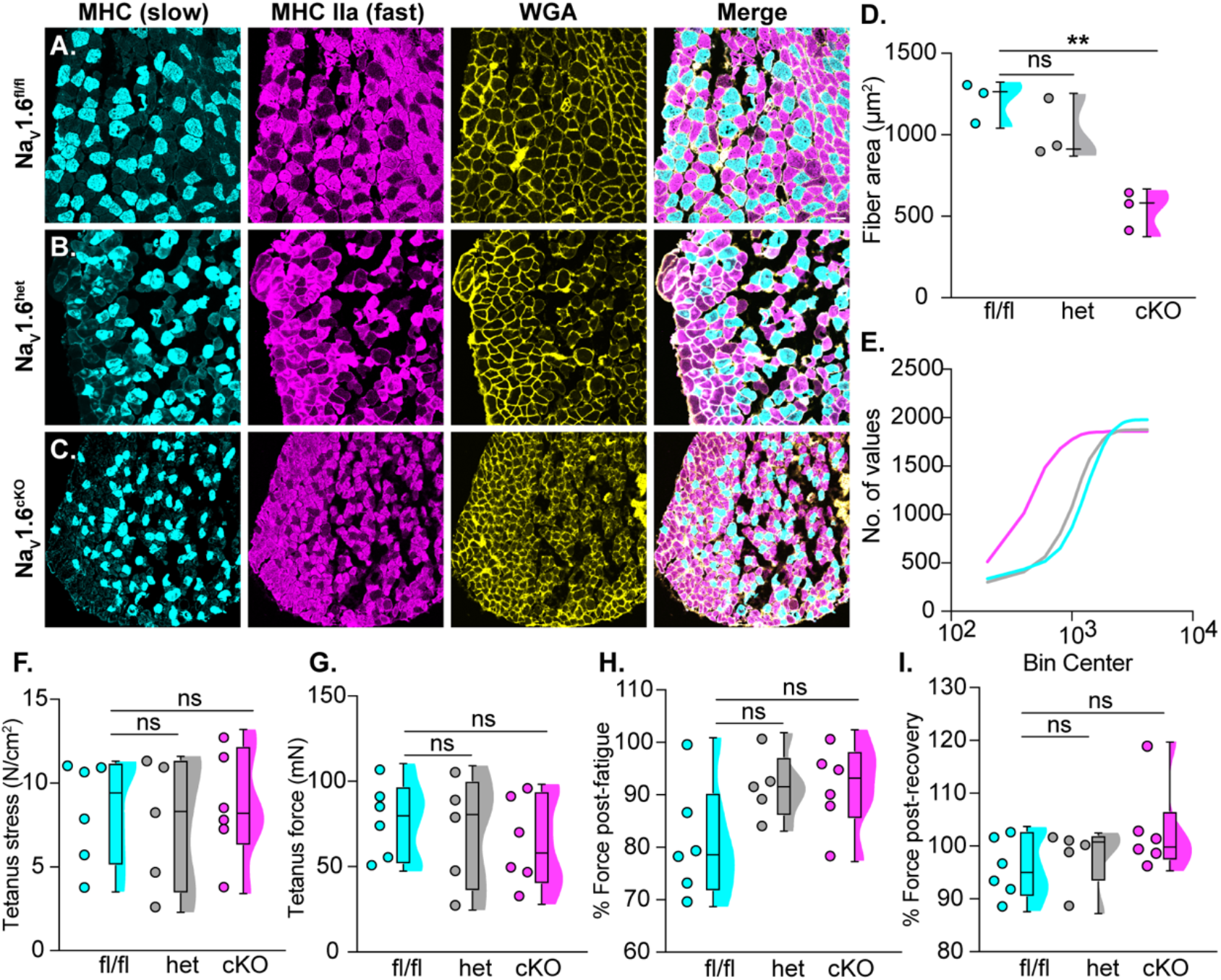
Loss of proprioceptive feedback alters skeletal muscle development in NaV1.6^cKO^ mice. Representative images of muscle fibers from the soleus of (**A**) NaV1.6^fl/fl^, (**B**) NaV1.6^het^, and (**C**) NaV1.6^cKO^ mice. Images were acquired with a 20x 0.75 NA air lens. Myosin heavy chain (MHC), labels slow twitch muscle fibers (cyan), MHC type IIa labels fast twitch muscle fibers (magenta), and wheat germ agglutinin (WGA, yellow) labels the cell membrane of muscle fibers. (**D** and **E**) Quantification of muscle fiber anatomy. (**D**) Fiber area, NaV1.6^het^ p = 0.337, NaV1.6^cKO^ p = 0.006, compared to NaV1.6^fl/fl^. (**E**) cumulative distribution plots showing the muscle fiber area in the soleus between NaV1.6^fl/fl^ (cyan), NaV1.6^het^ (grey), and NaV1.6^cKO^ (magenta) mice. (**F** to **I**) Quantification of intrinsic properties of soleus muscle. (**F**) Tetanus stress, NaV1.6^het^ p = 0.926, NaV1.6^cKO^ p = 0.990, compared to NaV1.6^fl/fl^. (**G**) Tetanus force, NaV1.6^het^ p = 0.925, NaV1.6^cKO^ p = 0.690, compared to NaV1.6^fl/fl^. (**H**) Percentage of force post-fatigue, NaV1.6^het^ p = 0.189, NaV1.6^cKO^ p = 0.150, compared to NaV1.6^fl/fl^. (**I**) Percentage of force post-recovery, NaV1.6^het^ p = 0.851, NaV1.6^cKO^ p = 0.293, compared to NaV1.6^fl/fl^. Each dot represents a single animal. Box and whisker plots represent maximum, minimum, median, upper and lower quartiles of data sets. A one-way ANOVA (Dunnett’s post-hoc comparison) was used to determine statistical significance. Scale bar=50 μm.

We next asked whether moderately impaired proprioceptive signaling due to loss of NaV1.1 had the same effect on skeletal muscle development or function. We examined muscle fiber composition of NaV1.1 mice of all genotypes and did not observe any significant changes in fiber area (Fig. 8 A to E). Additionally, the proportion of Type I and Type IIa fibers were similar between genotypes (Fig. S7 E to F). Functional analysis of intrinsic muscle properties in the soleus of NaV1.1^cKO^ mice did not differ compared to NaV1.1^het^ or NaV1.1^fl/fl^ mice (Fig. 8 F to I), suggesting moderately impaired proprioceptive feedback does not result in deficits in skeletal muscle developmental at the anatomical or functional levels. Taken together, these experiments unveil a novel role for proprioceptive feedback in skeletal muscle development.

**Fig. 8.**
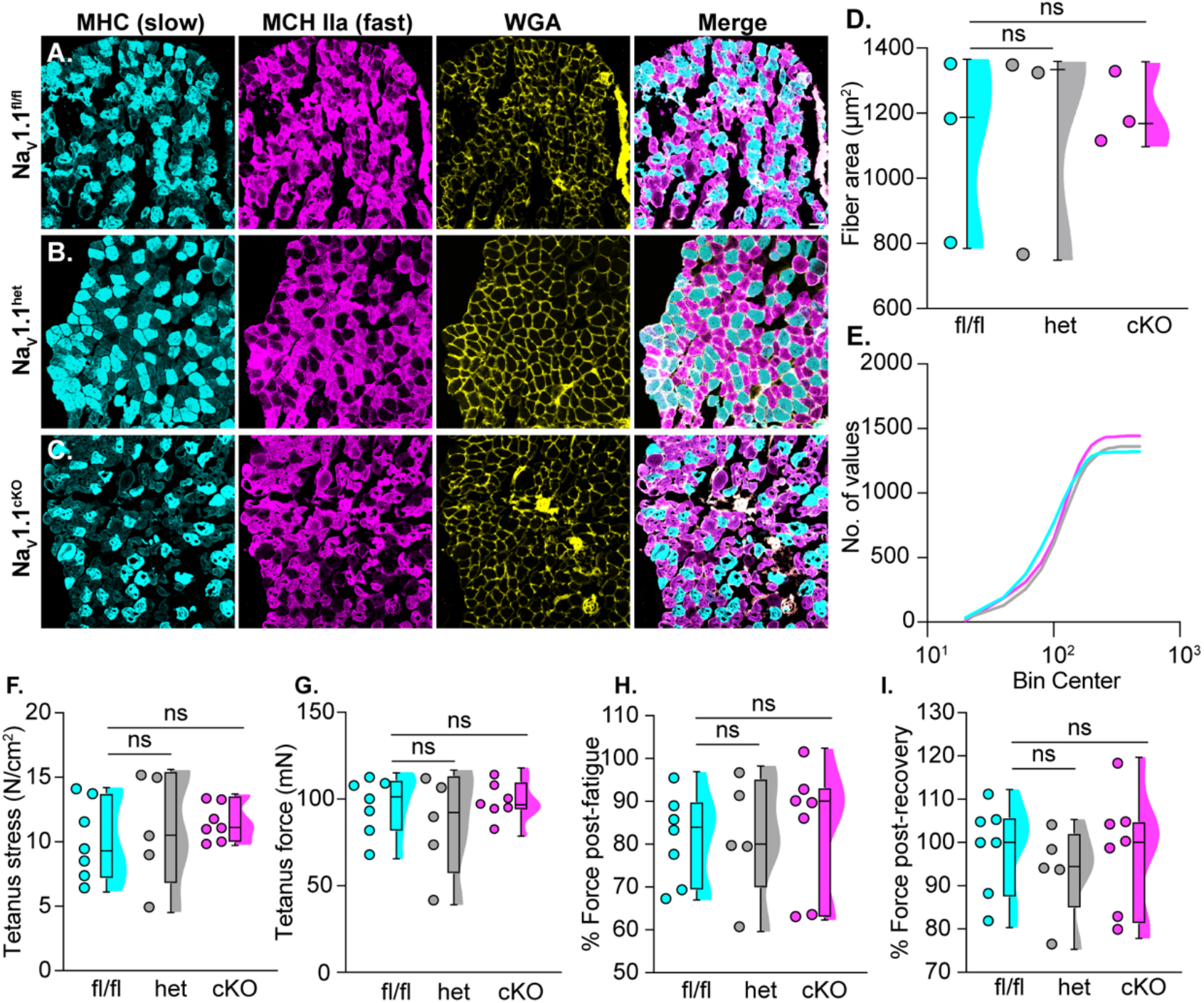
Mildly impaired proprioceptive feedback does not impair skeletal muscle development. Images of muscle fibers from (**A**) NaV1.1^fl/fl^, (**B**) NaV1.1^het^, and (**C**) NaV1.6^cKO^ soleus muscle. Images were acquired with a 20x 0.75 NA air lens. Myosin heavy chain (MHC) labels slow twitch muscle fibers (cyan), MHC type IIa labels fast twitch muscle fibers (magenta), and wheat germ agglutinin (WGA, yellow) labels the cell membrane of muscle fibers. (**D** and **E**) Quantification of muscle fiber anatomy. (**D**) Fiber area, NaV1.1^het^ p = 0.9827, NaV1.1^cKO^ p = 0.880, compared to NaV1.1^fl/fl^. (**E**) cumulative distribution plots showing the muscle fiber area between NaV1.1^fl/fl^ (cyan), NaV1.1^het^ (grey), and NaV1.1^cKO^ (magenta). (**F** to **I**) Quantification of intrinsic properties of soleus muscle. (**F**) Tetanus stress, NaV1.1^het^ p = 0.841, NaV1.1^cKO^ p = 0.596, compared to NaV1.1^fl/fl^. (**G**) Tetanus force, NaV1.1^het^ p = 0.624, NaV1.1^cKO^ p = 0.978, compared to NaV1.1^fl/fl^. (**H**) Percentage of force post-fatigue, NaV1.1^het^ p = 0.999, NaV1.1^cKO^ p = 0.934, compared to NaV1.1^fl/fl^. (**I**) Percentage of force post-recovery, NaV1.1^het^ p = 0.724, NaV1.1^cKO^ p = 0.993, compared to NaV1.1^fl/fl^. Each dot represents a single animal. Box and whisker plots represent maximum, minimum, median, upper and lower quartiles of data sets. A one-way ANOVA (Dunnett’s post-hoc comparison) was used to determine statistical significance. Scale bar=50 μm.

### NaV1.1 and NaV1.6 occupy distinct cellular domains within muscle spindles

Our results demonstrate that NaV1.1 and NaV1.6 have differential roles in proprioceptive signaling. We next sought out to define the mechanistic basis of their distinct and nonredundant contributions. At the biophysical level, NaV1.1 and NaV1.6 are functionally very similar, and both contribute to peak, persistent, and resurgent sodium currents (*26–28*). Previous studies examining central neurons, however, show that NaV1.1 and NaV1.6 occupy distinct excitable domains (*29–31*), suggesting that differences in cellular localization could dictate the unique roles these channels play in proprioception. Compared to the central nervous system, our understanding of NaV expression in sensory terminals is extremely poor. We therefore set out to examine the expression patterns of NaV1.1 and NaV1.6 in proprioceptive end organs. We focused our analysis on muscle spindles, as these structures comprise two of the three proprioceptor functional classes (*1*) and are the afferent endings from which we recorded in *ex vivo* muscle nerve experiments (Figs. 2 and 3). We first labeled for NaV1.6 channels and observed discrete, high density clusters across the spindle that resembled action potential initiation zones (Fig. 9A to C). We observed approximately 2 NaV1.6^+^ clusters per muscle spindle section (Fig. 9D), though this is likely an underestimation of the total number of clusters per entire spindle. In central neurons, NaV1.6 has been shown to play a major role in signal initiation and propagation due to its expression at the distal axon initial segment (AIS) and nodes of Ranvier (*29*, *31*, *32*). Thus, we co-labeled with the AIS marker Ankyrin-G (AnkG, Fig. 9 E to G, *33*) and found that 100% of NaV1.6 clusters colocalize with AnkG (Fig. 9 H). To determine if these clusters were *bona fide* heminodes or nodes of Ranvier, we co-labeled with the juxtaparanode marker CASPR (*34*). Indeed, triple immunolabelling experiments found NaV1.6 clusters flanked by two CASPR+ signals near VGLUT1^+^ muscle spindles (Fig. 9 I to L), as well as in myelinated axons of the sciatic nerve (Fig. S8). Furthermore, we also observed NaV1.6 channel clusters flanked by a single CASPR^+^ signal within muscle spindles, indicative of the presence of NaV1.6^+^ heminodes (Fig. 9 M). We ensured the specificity of NaV1.1 and NaV1.6 antibodies using tissue harvested from NaV1.1^cKO^ and NaV1.6^cKO^ mice, respectively (Fig. S9). These findings reveal that NaV1.6 is expressed only at heminodes within muscle spindles and nodes of Ranvier of proprioceptors, where it likely plays a direct role in signal initiation and propagation.

**Fig 9.**
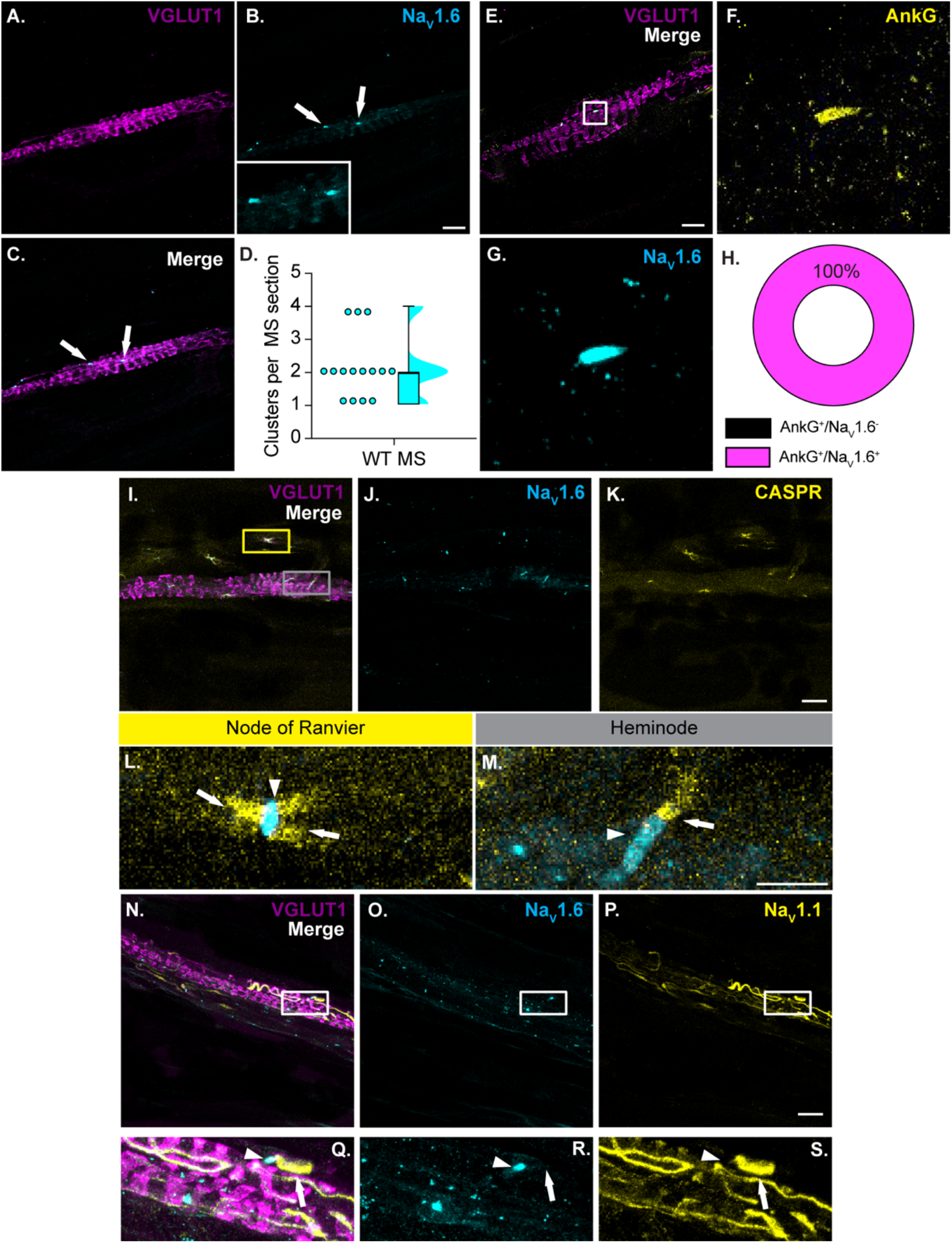
NaV1.1 and NaV1.6 localize to discrete cellular regions in muscle spindles. VGLUT1 (magenta, **A**) labeled muscle spindles express clusters of NaV1.6 (cyan, **B** and **C**). Arrows denote clusters of NaV1.6. (**D**) Quantification of the number of NaV1.6 clusters per muscle spindle (n=15 spindles). (**E** to **H**) NaV1.6 clusters (**G**) colocalize with Ankyrin-G (yellow, **E** and **F** are insets from G). (**H**) Quantification of the percentage of Ankyrin G clusters that colocalize with NaV1.6 clusters (n=9 spindles). (**I** to **M**) Co-labeling of NaV1.6 (**J**) with juxtaparanode maker CASPR (**K**, yellow) reveal proprioceptor nodes of Ranvier (**L**) and heminodes (**M**) (n=10 spindles). Nodes of Ranvier were identified by two CASPR+ signals (arrows) flanking NaV1.6 clusters (arrowhead). Heminodes were identified by a single CASPR+ signals juxtaposed to NaV1.6 cluster. (**N** to **S**) Co-labeling of NaV1.6 (**O**) with NaV1.1 (**P**, yellow) show discrete cellular expression patterns. (**Q** to **S**) Arrowhead denotes NaV1.6 channels and arrows denote NaV1.1 channels. N=3-5 mice. Inset scale bars=10 μm. Scale bars=20 μm

If NaV1.1 and NaV1.6 differentially regulate electrical signaling in proprioceptors through distinct cellular localization patterns, co-labeling for both ion channels should reveal non-overlapping expression patterns. In line with our hypothesis, we find a notable difference in NaV1.1 localization in muscle spindles compared to NaV1.6 (Fig 9 N and Q). In contrast to the discrete clusters of NaV1.6, we observe NaV1.1 localization is broader, but restricted to more equatorial wrappings within muscle spindles and in some presumptive axons entering the muscle spindle (Fig. 9 S).

Given we found developmentally dependent roles for NaV1.1 and NaV1.6 in proprioceptor synaptic transmission in the spinal cord (Figs. 4 and 5), we asked whether NaV localization is dynamic during postnatal development. We labeled for NaV1.1 and NaV1.6 in muscle spindles from the EDL of P7 and P14 C57Bl6/J mice (Fig. S10), the timepoints at which we observed a change in the requirement for either channel to motor function (Figs. S2 and S4). At P7, we observed no NaV1.6 clusters within spindles, though we did find some clusters near spindles, which could represent nodes of Ranvier. At P14, clusters of NaV1.6 begin to emerge within muscle spindles, though these clusters appear smaller and less frequently than those observed in adult muscle spindles. We detected little to no NaV1.1 immunoreactivity in muscle spindles at P7 and P14, suggesting either no or low expression of this channel at these timepoints. Given the significant change in functional grasping observed in NaV1.1^cKO^ at P14 (Fig. S4E) this would suggest NaV1.1 may serve a role in sensory transmission outside of the muscle spindle at this timepoint in postnatal development. Thus, we propose that the unique localization patterns of NaV1.1 and NaV1.6, which are dynamically regulated during postnatal development, confer the unique contributions of each channel to electrical signaling in proprioceptors.

## Discussion

The discovery of Piezo2 shared the 2021 Nobel Prize in Physiology and Medicine due to its essential role in the function of various mechanosensory neurons. In proprioceptors, Piezo2 initiates muscle mechanotransduction signaling (*4*, *5*); however, the downstream ion channels responsible for transmitting proprioceptive information to central circuits has remained mysterious. Here, we demonstrate that NaVs differentially encode mammalian proprioception, and we predict this is largely due to differences in channel localization within proprioceptors (Fig. 10). Our prior work found that NaV1.1 is essential for maintaining consistent and reliable proprioceptor encoding of static muscle stretch (*7*); this is consistent with its expression at sensory wrappings of muscle spindles where is it poised to amplify Piezo2-mediated mechanotransduction currents. Conversely, due to its localization at heminodes within muscle spindles and nodes of Ranvier, NaV1.6 has an obligate role in initiating proprioceptor action potential firing. This is in line with *ex vivo* recordings from NaV1.6^cKO^ afferents, in which responses to both dynamic and static muscle movement are abolished (Figs. 2 and 3). Furthermore, this demonstrates that the activity of Piezo2 and other ion channels in proprioceptors cannot compensate for the loss of NaV1.6. Thus, we conclude that NaV1.6 is equally essential for mammalian proprioception as Piezo2. To our knowledge, this is the first study to investigate and define unique and nonredundant roles for NaVs in somatosensory encoding.

**Fig. 10.**
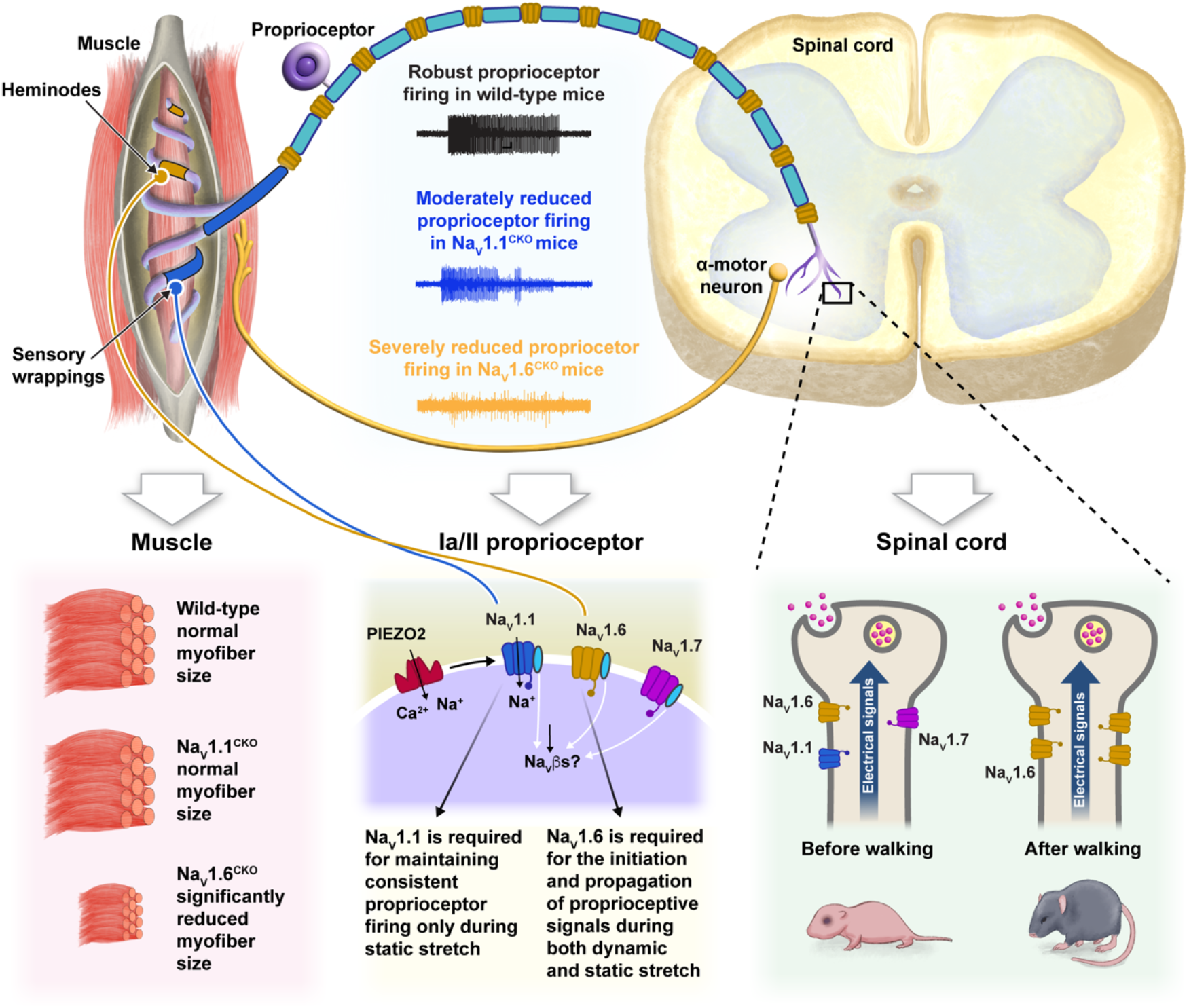
Model of proprioceptive transmission by NaV1.1 and NaV1.6. Upon muscle stretch, Piezo2 (red) transduces mechanical stimuli into electrical potentials. Following Piezo2 activation, NaV1.1 (blue) expressed in muscle spindle sensory terminals drives consistent proprioceptor firing during static muscle stretch. NaV1.6 (yellow) localized to heminodes and nodes of Ranvier initiate and propagate all proprioceptive signals from muscle spindles to spinal cord. It is likely prior to walking, there is functional redundancy of NaVs in proprioceptive axons. After walking behaviors emerge however, proprioceptive synaptic transmission is dependent on NaV1.6. Deletion of NaV1.6 in all sensory neurons led to a significant decrease in skeletal muscle fiber size that was not present in NaV1.1^cKO^ muscle, suggesting that complete loss of proprioceptive feedback non-cell autonomously regulates skeletal muscle development.

At the behavioral level, deletion of NaV1.1 or NaV1.6 in sensory neurons led to phenotypically distinct motor deficits. Previously, we reported that NaV1.1^cKO^ mice display uncontrollable intention-like tremors and poor motor coordination (*7*). Here we show that deletion of NaV1.6 in sensory neurons resulted in even more severe ataxia that precluded testing in the rotarod (Fig. 1). Notably, though more impaired than NaV1.1^cKO^ mice, NaV1.6^cKO^ mice did not display intention tremors. The distinct motor phenotypes that result from conditional deletion of NaV1.1 or NaV1.6 likely arise from differences in their respective contributions to proprioceptor function and development (Figs. 2, 3 and 6). It should be noted, however, that the behavioral phenotypes in these models cannot solely be attributed to proprioceptor dysfunction, as NaV1.1 and NaV1.6 are also expressed in tactile sensory neurons (*35*), which also contribute to motor behaviors (*36*, *37*). Nevertheless, at the afferent level, we show NaV1.6 is fundamentally essential for electrical signaling in proprioceptors, which contrasts the selective impairment on static stretch encoding observed in NaV1.1^cKO^ mice. Furthermore, NaV1.1 deletion had no significant impact on muscle spindle or skeletal muscle development; therefore, it is likely that the deficits observed in NaV1.1^cKO^ are purely electrical in nature. By contrast, in addition to loss of proprioceptive transmission, NaV1.6^cKO^ mice had abnormal muscle spindle structure (Fig. 6) and significantly reduced skeletal muscle fiber size (Fig. 7), which suggests that the severe motor deficits in NaV1.6^cKO^ may be caused by both cell-autonomous and non-cell-autonomous mechanisms. Interestingly, prior work on *Scn8a^med^* mice show that global inactivation of NaV1.6 resulted in a similar ataxic-like phenotype, as well as muscle atrophy and weakness (*12*, *22*). Our data show loss of NaV1.6 function in sensory neurons, particularly proprioceptors, are a major contributor to the motor and muscle impairments in *Scn8a^med^* mice. This may have broader implications for interpreting the clinical manifestations associated with disease causing *Scn8a* mutations, which often result in motor dysfunction (*11*).

We found that a single copy of NaV1.6 in sensory neurons was sufficient for normal motor function in adults (Fig. 1), despite being haploinsufficient in *ex vivo* muscle nerve recordings in response to vibration (Figs. 3). Indeed, in response to vibratory stimuli, afferents from NaV1.6^het^ animals were less likely to entrain to sinusoidal vibration, particularly at 25μm stimulus amplitudes (Fig. 3 B and D). Why these functional deficits in NaV1.6^het^ afferents do not manifest at the behavioral level is unclear. It should be noted that at P14 we observed NaV1.6 haploinsufficiency in *ex viv*o monosynaptic reflex recordings (Fig. 4 and Table S1) and in a motor coordination assay (Fig S2).

It is therefore possible that compensatory mechanisms in NaV1.6^het^ animals come into play in early adulthood. Another possibility is that NaV1.6^het^ animals possess more subtle motor deficits that were unresolvable in the open field and rotarod. More sensitive kinematic analyses with higher spatial and temporal resolution will be required to investigate the extent of NaV1.6 haploinsufficiency for proprioceptor-driven motor behaviors.

We have very limited knowledge about the localization of ion channels within somatosensory end organs. This information is important for understanding how electrical signals arise within structurally complex sensory terminals, which can be damaged during pathological conditions or aging (*1*, *38–40*). We show that NaV1.1 and NaV1.6 occupy distinct cellular compartments within proprioceptive muscle spindle end organs, which we predict underlies their differential roles in encoding proprioceptive signals. While differences in biophysical properties could also underly the differential roles of NaV1.1 and NaV1.6 to proprioceptive transmission, these channels share many functional similarities. They both rapidly activate and inactivate, can generate peak, persistent and resurgent currents, and have similar recovery from inactivation kinetics (*7*, *27*, *28*, *41–43*). Prior work in neurons of the central nervous system is consistent with our hypothesis that localization dictates NaV contributions to proprioceptor function. For example, in retinal ganglion cells and motor neurons, NaV1.6 is preferentially expressed at the distal axon initial segment, suggesting a primary role in signal initiation (*44*, *45*). Furthermore, studies have shown that NaV1.6 is the dominant isoform at nodes of Ranvier, playing a key role in action potential propagation in myelinated axons (*31*, *45*). In contrast, NaV1.1 is localized to the soma and proximal AIS, where it aids in repetitive firing in fast spiking neurons of the brain (*14*, *46*). This is in line with our model whereby NaV1.1 amplifies mechanotransduction currents from Piezo2 to maintain sustained action potential firing during static stretch.

Interestingly, we found that 100% of NaV1.6 immunoreactivity colocalizes with AnkG. AnkG is known to anchor NaVs within the AIS of central neurons (*32*, *33*, *47*, *48*); thus, our findings indicate muscle spindles possess several NaV1.6-expressing action potential initiation zones. Surprisingly, we never observed broad AnkG immunoreactivity in muscle spindle sensory wrappings, indicating AnkG does not colocalize with NaV1.1, despite its known colocalization with NaV1.1 at the AIS in other neurons of the central nervous system (*30*, *49*). The mechanisms that anchor NaV1.1 to sensory terminals remain unknown. Scaffolding proteins known to colocalize with NaV1.1 include βIV-spectrin, auxiliary NaVβ subunits, and fibroblast growth factors (*14*, *50*, *51*). Proximity proteomic approaches could identify specific molecular players involved in NaV1.1 channel organization within proprioceptive end organs.

Another surprising result from our study was the developmentally dependent manner in which NaVs contribute to proprioceptor synaptic transmission in the spinal cord (Fig. 10). Ventral root recordings from mice at ages P6 to P11 revealed that neither NaV1.1 or NaV1.6 are required for the proprioceptor-mediated monosynaptic reflex response at this age. Conversely, by P14, when proprioceptors are nearing molecular maturation and weight bearing locomotion has emerged, NaV1.6 becomes absolutely critical for this circuit. There are two principal interpretations for these data. First, prior to walking behaviors, neither NaV1.1 or NaV1.6 contribute to proprioceptor synaptic transmission onto motor neurons. This interpretation would be consistent with previous studies in myelinated neurons of the retina, whereby the onset of eye opening corresponds with a developmental switch from NaV1.2 to NaV1.6 (*44*). It is possible that another NaV subtype, such as NaV1.7, is the dominant channel in early postnatal development. Alternatively, another interpretation is that in early postnatal development, there is functional redundancy among NaV subtypes, and loss of one is insufficient to impair synaptic transmission. This is line with a previous study that found the presence of multiple NaV isoforms in sensory axons as early as P7 (*52*). We favor the latter interpretation because functional redundancy is a common phenomenon in the developing nervous system, and NaV1.7 does not appear to play a significant role in mammalian proprioception. Nevertheless, both interpretations indicate that the cellular trafficking mechanisms governing the stability of each NaV subtype in this circuit are independent of one another, as loss of NaV1.6 did not result in compensation by other NaVs following the onset of weight bearing locomotion. Notably, our analyses of the monosynaptic reflex are consistent with behavioral analyses carried out in P7 and P14 mice, where motor function in P7 NaV1.1^cKO^ and NaV1.6^cKO^ mice was largely intact but declined by P14 (Figs. S2 and S4). Interestingly, at P7 we observed little-to-no immunoreactivity of NaV1.1 or NaV1.6 in muscle spindles (Fig. S9). By P14, NaV1.6 clusters begin to appear within the spindle, while NaV1.1 immunoreactivity remained weak. This suggests temporally distinct regulation of NaV localization at peripheral end organs compared to central circuits.

We find that loss of NaV1.6, but not NaV1.1, resulted in disrupted muscle spindle development. This is in line with recent findings that show mechanosensory neuron end organ development is activity dependent (*21*). Interestingly, previous work found deletion of Piezo2 in proprioceptors (Piezo2^cKO^) did not alter muscle spindle structure (*5*); however, these experiments were carried out in 4–5- week-old mice, whereas our analysis of muscle spindle structure was carried out in mice ages 8-12 weeks. This raises the possibility that electrical activity is required for the maintenance, but not development, of muscle spindle structure.

Surprisingly, NaV1.6^cKO^ mice showed significantly reduced skeletal muscle fiber size, highlighting a potential non-cell-autonomous role for proprioceptive feedback in skeletal muscle development or maintenance (Figs. 7 and 10). This effect was not seen in NaV1.1^cKO^ mice, but is consistent with a study that found deletion of Piezo2 in proprioceptors led to non-cell-autonomous deficits in spine alignment and hip joint formation (*53*). Despite the smaller size of NaV1.6^cKO^ skeletal muscle fibers, all NaV1.6^cKO^ intrinsic muscle properties were similar to NaV1.6^het^ and NaV1.6^fl/fl^ mice. Grip strength in NaV1.6^cKO^ mice, however, was significantly weaker compared to other genotypes. These findings suggest that reduced grip strength in NaV1.6^cKO^ mice is likely due to impaired motor neuron activation of skeletal muscle and not due to changes in intrinsic muscle function. Interestingly, prior studies have identified dysfunction in proprioceptive spinal cord circuits in mouse models of spinal muscular atrophy and amyotrophic lateral sclerosis (*19*, *54–56*). An intriguing possibility is that loss of proprioceptive feedback onto motor neurons in NaV1.6^cKO^ mice (Fig. 4) could lead to pathophysiological phenotypes similar to those observed in neuromuscular or motor neuron disease.

Mutations in the genes that encode NaV1.1 and NaV1.6, *Scn1a* and *Scn8a*, respectively, are strongly associated with neurological diseases in which ataxia and motor developmental delays are prominent clinical manifestations (*12*, *26*, *46*). How proprioceptor dysfunction contributes to these disorders is unknown. The majority of Dravet syndrome patients have complete loss-of-function (LOF) of one copy of *Scn1a*; conversely, patient reported mutational variants in *Scn8a* are predominately gain-of-function (GOF). Our previous work found that proprioceptor afferents from NaV1.1^het^ mice also had impaired encoding static muscle stretch (*7*), suggesting that motor dysfunction in patients missing one functional copy of *Scn1a* could be in part sensory in nature; however, it is unclear how *Scn8a* GOF mutations would affect proprioceptor function. Patients harboring either *Scn1a* LOF and *Scn8a* GOF mutations have similar motor deficits (*11*); thus, one possibility is that *Scn8a* GOF mutations lead to use-dependent block of action potential firing in proprioceptors, which could contribute to the similar motor phenotypes observed in these different patient populations (*11*). Interestingly, there are a few reported cases of LOF mutations in *Scn8a* that lead to general ataxia (*57*), and our data suggest that proprioceptor dysfunction could contribute to their motor deficits.

In addition to NaV1.1 and NaV1.6, proprioceptors also express NaV1.7 (*7*). Mice and humans that lack NaV1.7 are insensitive to pain but do not exhibit prominent deficits in motor function, which suggests a limited role of NaV1.7 in mammalian proprioception (*9*, *10*). Alternatively, NaV1.1 and NaV1.6 may compensate for the developmental loss of NaV1.7, which occurs in constitutive genetic mouse models and human patients. It is possible that acute deletion of NaV1.7 in proprioceptors could reveal a previously overlooked contribution of the channel to proprioceptor function. Indeed, we found that NaV1.7 channels contribute to roughly one third of the somal whole-cell sodium current in genetically identified proprioceptors (*7*). Thus, a role for NaV1.7 in mammalian proprioception remains enigmatic.

A current limitation of the present study is the use of a sensory-neuron wide genetic targeting strategy, which makes interpretation of motor behavior and skeletal muscle impairments confounded by the loss of NaV1.1 or NaV1.6 in other sensory neuron populations, namely touch receptors. As mentioned above, deletion of NaV1.1 or NaV1.6 selectively in proprioceptors is not possible with currently available genetic tools, as the access point for Cre-driven deletion, parvalbumin, is also expressed in neurons of brain and spinal cord that are important for motor function. Despite this limitation, the direct role of NaV1.1 and NaV1.6 in proprioceptors was examined at the functional level in *ex vivo* muscle nerve recordings and spinal cord electrophysiology experiments, as well as in the structural analysis of VGLUT1 identified muscle spindles. To investigate the role of NaV channels with spatial precision, future experiments will require intersectional strategies for selective gene manipulations in proprioceptors.

Our data demonstrate that NaV1.1 and NaV1.6 play distinct and nonredundant roles in mammalian proprioception. This work is the first to define how NaVs uniquely shape somatosensory transmission and is also the first to show that NaVs occupy distinct cellular compartments in sensory neuron end organs. We predict our results are broadly applicable to other sensory neuron populations, namely mechanoreceptors, which also co-express NaV1.1 and NaV1.6. Furthermore, these data have important translational implications for understanding the motor deficits associated with NaV1.1 and NaV1.6 channelopathies.

## Materials and Methods

### Experimental design

#### Animals

Pirt^cre^ and Scn8a^fl/fl^ mice were a gift from Drs. Xinzhong Dong (Johns Hopkins University, *58*) and Miriam Meisler (University of Michigan, *59*), respectively. Scn1a^fl/fl^ (stock #041829-UCD) were purchased from the UC Davis MMRRC. All mice used are a C57BL/6J background (non-congenic). Genotyping was outsourced to Transnetyx. Animals use was conducted according to guidelines from the National Institutes of Health’s Guide for the Care and Use of Laboratory Animals and was approved by the Institutional Animal Care and Use Committee of UC Davis (#23049) and San Jose State University (#990, *ex vivo* muscle recordings.) NaV1.6^cKO^ mice were provided with wet food and hydrogels daily. Mice were maintained on a 12hr light/dark cycle, and food and water were provided ad libitum.

#### Animal behavior

Motor function was tested using three assays: rotarod, open-field, and grip strength. Behavioral assays were conducted from least to most invasive in the order of open field, grip strength, and rotarod. All behavioral assays were conducted between 8-10 weeks of age and the experimenter was blind to genotype. For the open field, mice were acclimated to the behavior room for 1 h prior testing. The open field apparatus consisted of a white square box with dimensions of 15x15x20 inches. A camera suspended above the open field tracked animal movement for a single 10-minute period using ANY-maze software. Following testing in the open field, mice were transported to a separate behavior room and allowed to acclimate for 1 h before being assayed in the grip strength and rotarod tests. A grip strength apparatus (IITC Life sciences, Woodland Hills, CA) with a metal grate was used. Mice held from the tail were placed on the metal grate and pulled horizontally away from apparatus once all four paws touched the grate. Mice were assayed across 6 trials with 5-minute intervals between trials. A rotarod machine (IITC) that has an accelerating rotating cylinder was used. NaV1.6^cKO^ mice were excluded from rotarod testing due to severe motor coordination deficits that prevented them from maintaining balance on the cylinder even in the absence of cylinder rotation. The averages of three trials across three consecutive training days were recorded.

#### *Ex vivo* muscle nerve recordings

Detailed methods on *ex vivo* muscle nerve recordings can be found in Wilkinson et al. 2012 (*16*). Briefly, extensor digitorum muscle and innervating peroneal branch of the sciatic nerve were dissected from adult mice and placed in a tissue bath of oxygenated Synthetic Interstitial Fluid at 24°C. Tendons were tied to a fixed post and lever arm of a dual force and length controller and transducer (300C-LR, Aurora Scientific, Inc.). The cut end of the nerve was suctioned into a bipolar glass electrode and connected to an extracellular amplifier with headstage (Model 1800, A-M Systems). Muscles were held at the length of maximal twitch contraction, Lo. For static stretch experiments, nine 4 s ramp-and-hold stretches were given at 2.5, 5, and 7.5% of Lo (Ramp speed was 40% Lo/s). Stretch lengths were repeated three times. For sinusoidal stimuli, sixteen 9 s sinusoidal vibrations were given at 5, 25, 50, and 100 μm amplitudes at varying frequencies (10, 25, 50, and 100 Hz). A rest period of 1 min was given between each length change. Resting discharge was quantified as the firing rate 10 s before stretch. Firing rate during the static phase of stretch was calculated 3.25-3.75 sec into the hold phase of stretch (LabChart Software, ADInstruments). The consistency of firing during static muscle stretch was found by calculating the interspike interval coefficient of variation during the plateau phase of stretch (CV = Std Dev/Mean of ISI 1.5 – 3.5s after ramp up). For dynamic responses, the average firing rates during the 9 s vibration was determined. Entrainment was defined as whether a unit could entrain in a 1:1 fashion to vibration stimulus. In most afferents we confirmed that they were Group Ia or II afferents by looking for a pause in firing during the shortening phase of contraction (a train of 60 stimulations of 0.5 ms pulse width were given at 1 Hz frequency from a 701C stimulator (Aurora Scientific)).

#### Spinal cord electrophysiology

Spinal cords were harvested from postnatal mice spanning the age-groups P6-11 and P14-18. The mice were deeply anesthetized with isoflurane, decapitated and eviscerated. We followed the protocol that has been used to record motor activity from mice of weight-bearing age using *ex vivo* spinal cord preparations (*60*, *61*). In brief, after evisceration, the preparation was pinned to a dissecting chamber and continuously perfused with ice-cold solution, comprising (in mM): 188 sucrose, 25 NaCl, 1.9 KCl, 10 MgSO4, 0.5 NaH2PO4, 26 NaHCO3, 1.2 NaH2PO4, 25 glucose, bubbled with 95 % O2/ 5 % CO2. The spinal cord was exposed following a ventral laminectomy and transected at the thoracic levels (T5-T8). The dorsal and ventral roots were isolated over the sixth lumbar segment, bilaterally, just proximal to the dorsal root ganglion. All other dorsal and ventral roots were trimmed, and the entire cord was removed from the vertebral column together with the attached roots and transferred to the recording chamber and continuously superfused with artificial cerebrospinal fluid (aCSF; concentrations in mM): 128 NaCl, 4 KCl, 1.5 CaCl2, 1 MgSO4, 0.5 NaH2PO4, 21 NaHCO3, 30 D-glucose) bubbled with 95 % O2 – 5 % CO2. A midsagittal hemisection was performed and the spinal hemicords were allowed to equilibrate in aCSF maintained at ambient temperature. After spinal cord isolation, dorsal and ventral roots at the sixth lumbar segment (L6) were placed into suction electrodes. The dorsal root was stimulated with single pulse stimulus, delivered every 30s, over 10 trials. The stimulus was delivered using a stimulus isolator unit (A365, World Precision Instruments) with current pulse amplitudes set at twice the threshold intensity of stimulation (2T, 0.1 ms pulse-width). Extracellular recordings were made at the ventral roots, the signal was filtered between 0.1–5000 Hz, amplified 1000 times (Model 1700, A-M Systems), digitized at 10 kHz using Digidata 1440A, acquired using Clampex software (v11.2, Molecular Devices), and saved on a computer for offline analysis. Stretch reflex parameters were extracted from the signals for each experiment, after averaging over the 10 trials, using Clampfit (v11.2, Molecular Devices).

#### Muscle Mechanics

Soleus muscles were prepared for *ex vivo* passive mechanical testing as previously described (*62*). Briefly, 7-0 sutures were cinched at the muscle-tendon of the soleus and EDL muscles. Suture loops were placed on hooks connected to the 300C-LR-Dual-Mode motor arm and force transducer (Aurora Scientific) such that the muscle remained within 28°C oxygenated Ringer’s solution. Twitches were induced using a 701C stimulator (Aurora Scientific) across a range of muscle lengths to determine the optimal length for isometric force generation (Lo). The Lo length corresponded to the length between the sutures on either muscle-tendon junction, as measured by calipers. Physiological cross-sectional area (PCSA) was calculated using the muscle length (Lm), mass (m), ratio of fiber length to Lo (Lf/Lo) and standard density of muscle (ρ=1.06 g/cm^3^; PCSA = m/Lo*(Lf/Lo)*ρ, (*63*).

Soleus muscles were subjected to active mechanical testing, which consisted of a series of 24 maximum isometric tetani (300 mA, 0.3 ms pulse width, 80 Hz pulse frequency, 800 ms pulse train) with 6 seconds of recovery in between each tetanus. Muscles were then given 300 seconds to recover before a final tetanus with the same parameters. Maximum isometric force was measured during the first, penultimate, and final tetanus protocol. Maximum forces were normalized to PCSA to give isometric specific tension. The highest isometric specific tension measured during each active protocol was reported as the isometric specific tension for each muscle. The percent of force maintained at the penultimate tetanus compared to the initial tetanus was recorded as the percent force post-fatigue. The percent of force maintained at the final tetanus compared to the initial tetanus was recorded as the percent force post-recovery. After active mechanical testing was completed, muscles were removed from the mechanical testing equipment, embedded in OCT, and flash frozen in liquid nitrogen cooled isopentane. Muscles were stored at -70°C until cryosectioning.

#### Tissue Processing

For muscle spindle immunolabeling experiments, mice were anesthetized using a ketamine/xylazine cocktail and transcardially perfused with PBS followed by 1% PFA. Extensor digitorum longus (EDL) muscle was then dissected in PBS and post-fixed for 30 min then washed in PBS before incubation in 30% sucrose solution overnight at 4C. Following cryoprotection, muscles were embedded in optimal cutting temperature (Fisher #4585) and stored in -80C until sectioning.

#### Immunohistochemistry

For immunolabeling experiments in muscle spindles, EDL muscles were sectioned (30μm) along the longitudinal axis. Tissue was incubated in blocking solution (0.1% PBS-T/5% normal goat serum in PBS) and the following primary antibodies were used: guinea pig anti-VGLUT1 (1:8000, Zuckerman Institute 1705) and rabbit anti-βIII tubulin (1:3000, Abcam #ab18207). Secondaries are as follows: anti-guinea pig 488 (1:1000, Thermo Fisher, A11073) and anti-rabbit 647 (1:1000, A32733). For muscle fiber typing experiments, soleus cross sections (20μm) were blocked in a solution containing 5% BSA in PBS. The following primary antibodies were diluted and incubated on muscle sections overnight: mouse IgG2b anti-myosin heavy chain type I (1:250, BA-F8, DHSB) and mouse IgG1 anti-myosin heavy chain type IIa (1:250, SC-71, DHSB). Slides were washed in PBS and the following secondaries were diluted in 2% BSA and incubated for 60 minutes: goat anti-mouse IgG2b 488 (1:500, A21141) goat anti-mouse IgG1 555 (1:500 A21127). After secondary antibody incubation slides were washed in PBS and mounted with FluoromountG with DAPI (SouthernBiotech 0100-20). For immunolabeling of NaVs the following primary antibodies were used: guinea pig anti-VGLUT1 (1:8000, Zuckerman Institute 1705), rabbit anti-NaV1.1 (2ug/ml, Neuromab A11954), mouse IgG1 anti-NaV1.6, mouse (6ug/ml, Neuromab K87A/10.2), rabbit anti-NaV1.6 (1:750, Alomone, ASC009), IgG2a anti-Ankyrin G (6ug/ml, N106/36.1), and chicken anti-neurofilament heavy (1:3000, Abcam ab4680). Secondaries used were as follows: anti- guinea pig 488 (1:1000, Thermo Fisher, A11073), mouse anti-IgG1 555 (1:1000, A21137), mouse anti IgG2a-647 (1:1000, A21240), anti-chicken 594 (1:1000, WA316328). All specimens were imaged in three dimensions on either a Zeiss LSM880 Airsyscan (63x oil objective, 1.4 NA) or Olympus LV3000 (60x oil objective, 1.4 NA) confocal microscope. Images were analyzed using ImageJ software.

#### Analysis of muscle spindle structure

Disruptions in muscle spindle sensory endings were quantified by colocalizing VGLUT1 immunoreactivity with DAPI to calculate a wrapping efficiency index (WEI). Intrafusal muscle fibers are identifiable in skeletal muscle based on mono- and bi-nucleation via DAPI^+^ staining. To analyze muscle spindle sensory wrappings without bias, we only analyzed sensory wrappings VGLUT1 sensory wrappings that overlapped with DAPI labeling. For example, in ImageJ, regions of interest (ROIs) were drawn around each sensory wrapping. If the ROI did not overlap with DAPI, it was not counted as a sensory wrapping and was excluded from the analysis. The total number of sensory wrappings (n) were counted and normalized to muscle spindle length (l). The following equation was used to calculate the WEI:

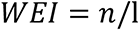

#### Experimental Design and statistical analysis

Summary data are presented as mean ± SD, from n cells, afferents, sections, or N animals. All analysis of immunofluorescent images contained at least 3 biological replicates per condition. Investigator was blinded to genotypes during analysis. For all behavioral, electrophysiological, and mechanics experiments the investigator was blind to genotype. To determine differences in entrainment properties between NaV1.6^fl/fl^, NaV1.6^het^, and NaV1.6^cKO^ we used a logistic regression analysis (SPSS), is a statistical model that calculates the log-odds of an event (i.e. entrainment or non-entrainment) as a linear combination of one or more independent variables (vibration frequency, vibration amplitude and genotype). All other statistical testing was carried out using Prism 10.1 (Graphpad software). Statistical differences were determined using parametric tests for normally distributed data and nonparametric tests for data that did not conform to Gaussian distributions or had different variances. Statistical significance in each case is denoted as follows: *p<0.05, **p<0.01, ***p<0.001, and ****p<0.0001. Source files for each figure can be found on Mendeley.

## Acknowledgments

We would like to thank Drs. Xinzhong Dong and Miram Meisler for sharing mouse lines, and Dr. James Trimmer for providing support and guidance on immunolabeling experiments. Thanks to Griffith and Contreras Lab members for helpful discussions. Additional support was provided by. Core facilities were supported by P30 EY12576.

## Author contributions

Conceptualization: CME, TNG

Methodology: CME, CN, SO, JRD, ARM, YM, ELM, SG, RPW, SEB, LS, KW, TNG

Investigation: CME, CN, SO, JRD, ARM, YM, ELM, SG, RPW, SEB

Visualization: CME, TNG Supervision: CME,TNG Writing—original draft: CME

Writing—review & editing: CME, TNG

## Data and materials availability

All data are available in the main text or the supplementary materials. Source data for each figure can be found on Mendeley.

## Supplementary Materials

### Supplementary Methods

#### Assessment of motor function during at postnatal day 7 and 14

Floxed, heterozygous, and conditional knockout P7 and P14 pups from the NaV1.1 and NaV1.6 mouse lines were assayed for motor dysfunction. The experimenter was blind to genotype.

#### Behavioral assays at postnatal day 7

Hindlimb foot angle: In a clear empty mouse cage, a camera was positioned from below and above to record the pup as it moved around the cage. The pup was gently prodded by touching its tail to motivate the pup to move. An average of three pictures were taken from above and below. Using the acquired pictures, measures of the foot angle of the pups were performed using Fiji ImageJ software by drawing a line from the end of the heel/shin to the tip of the middle toe on each hindlimb and measuring the angle of the intersecting lines. Measurements were only taken when the pup was performing a full stride in a straight line and both feet were flat on the ground. Three to five sets of foot angles were measured per pup and used to calculate the average angle for each pup tested.

Righting reflex: Pups were placed on their backs on a bench pad and held in that position for 5 seconds. The pups were released, and the time it took for the pup to return to the prone position was recorded. This was repeated for a total of three trials and the average righting reflex latency was calculated. Intertrial rest periods were 60 seconds.

Hindlimb strength: Using a 50ml conical tube with cotton ball padding at the bottom, the pup was gently placed face down into the tube with its hind limbs hung over the rim. The latency for the pup to fall into the tube was recorded. The test was ended at 60 seconds if the pup did not fall. Each pup was only tested one time to avoid exhaustion.

Grasping reflex: Pups were held by the scruff of the neck in a similar way to the how it is carried by the mother. The pad of each individual mouse paw was stroked using the wooden stick of a cotton tip applicator. The grasp reflex was determined present if the mouse paw curled around the wooden stick. Mice received a score of zero if all four paws had the grasp reflex present, a score of 1 if one paw did not have the grasp reflex present, a score of 2 if two paws did not have the grasp reflex present, and score of 3 if three paws did not have the grasp reflex present, and a score of 4 if four paws did not have the grasp reflex present.

#### Behavioral assay at postnatal day 14

Modified limb coordination assay: This test was used to determine differences in grip, balance and limb coordination at postnatal day 14. Pups were placed on a wire grid with metal poles running parallel to each other, approximately 8 millimeters apart with a diameter of 3 millimeters. The pups were left on the grid for five seconds and were scored by their ability to grip and balance with each individual limb without the paw slipping in between the metal bars. Pups that could grip/balance with all four limbs received a score of 0, pups that could grip/balance with only three limbs received a score of 1, pups that could grip/balance with only two limbs received a score of 2, Pups that could grip/balance with only one limb received a score of 3, and pups that could grip/balance with none of their limbs received a score of 4. The assay was repeated a total of three times with 30 seconds in between tests, and the average of the three trials was reported.

#### Multiplex *in situ* hybridization

DRG were harvested from adult (10-15 week-old) Pirt^Cre^;NaV1.6^fl/fl^ mice of both sexes. DRG was sectioned at 25 μm sections and were processed for RNA *in situ* detection using a modified version of manufacture protocol (Advanced Cell Diagnostics) as previously described (Griffith 2019 and Espino 2022). The following probes were used: Pvalb (421931C1, mouse) and Runx3 (451271-C3, mouse). Following in situ hybridization, sections were incubated in blocking solution (5% NGS, 0.1% PBS-T) for 1hr at room temperature (RT). Tissue was incubated in rabbit βIII-Tubulin primary antibodies (1:3000, Abcam ab41489) overnight at 4°C overnight. Tissue was treated with anti-rabbit 594 (1:1000, Invitrogen, A11037) secondary antibodies for 30 min at RT. Sections were mounted with Fluoromount-G with DAPI and imaged in three dimensions on Olympus confocal (FV3000) using 40x 0.90 NA water objective lens. Images were analyzed using ImageJ software.

#### Immunolabeling of muscle spindles in postnatal development

Immunohistochemistry of EDL harvested from P7 and P14 C57Bl6/J mice was performed. EDL was sectioned (25 μm) on a cryostat and sections were labeled using the following primary antibodies: rabbit polyclonal NaV1.1 (2μL/mL, Neuromab, NACH AP11954), guinea pig anti- VGLUT1 (1:8000, Zuckerman Institute, 1705), and chicken anti-NFH (1:3000, Abcam, ab4680), rabbit polyclonal anti- NaV1.6 (1:750, Alomone Labs, ASC-009). Secondary antibodies used were as follows: anti-rabbit 594 (1:500, Thermo Fisher, A32740), anti-guinea pig 647 (1:1000, Thermo Fisher, A11073), and anti-chicken 647 (Thermo Fisher, A32733), anti-mouse IgG2a 555 (A21137). Specimens were mounted with Fluoromount-G with DAPI (SouthernBiotech, 0100-20). All specimens were imaged in three dimensions on an Olympus FV3000 confocal microscope using 60x NA 1.4 oil objective lens. Images were analyzed using ImageJ software.

## Supplementary figure and legends

**Fig. S1.**
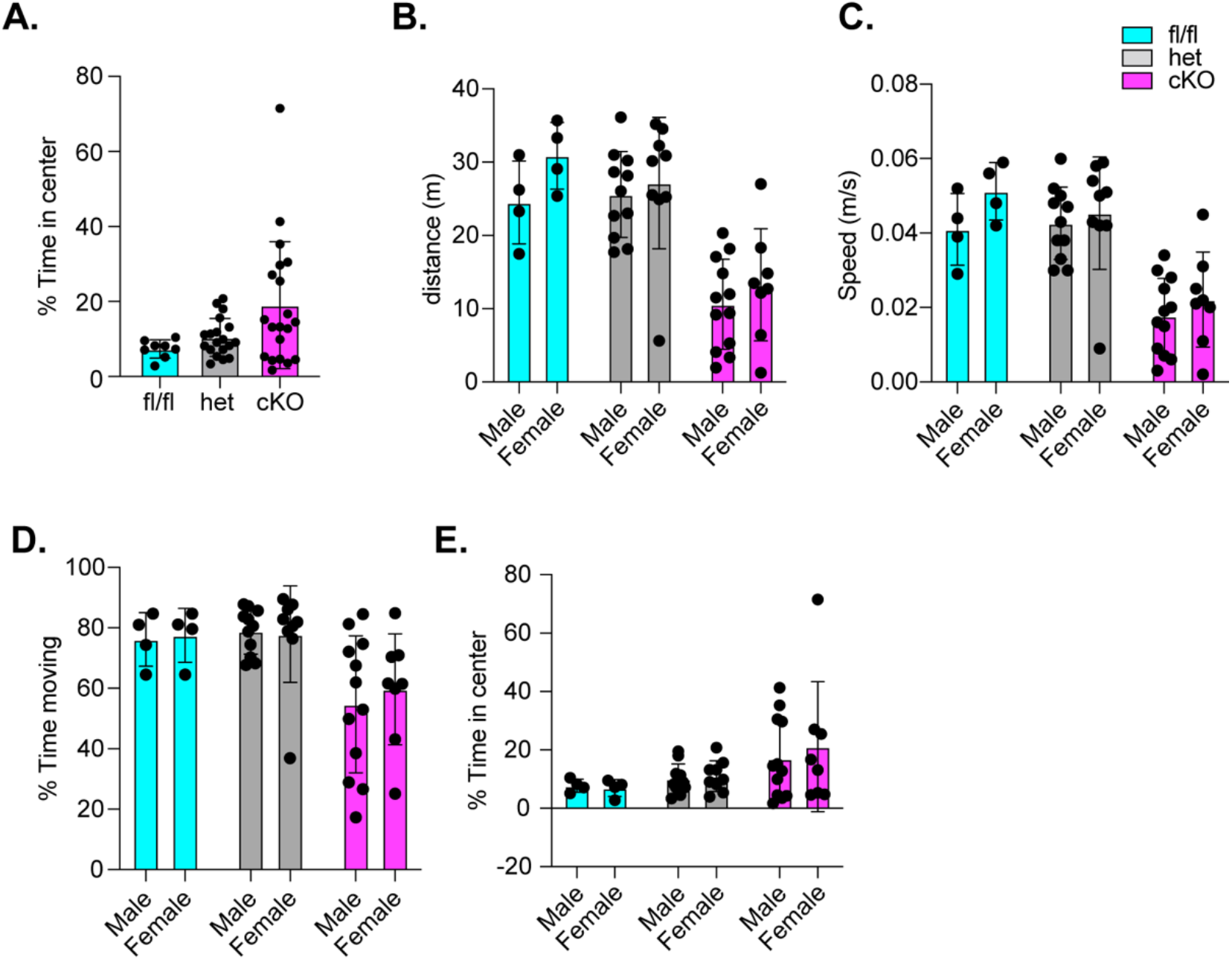
Motor behavior analysis of mice lacking NaV1.6 in sensory neurons. (**A**) Quantification of percent time spent in the center during a 10 min open-field trial. A Kruskal- Wallis test with Dunn’s Post-hoc comparison was used to determine statistical significance. NaV1.6^het^ p=0.4206, NaV1.6^cKO^ p=0.1026 compared to NaV1.6^fl/fl^ . NaV1.6^fl/fl^ n = 8, NaV1.6^het^ n=20, NaV1.6^cKO^ n =20. No significant differences were observed for motor behaviors male and female mice of all genotypes. (**B**) Distance moved. (**C**) Speed. (**D**) Percent time moving. (**E**) Percent time spend in center.

**Fig. S2.**
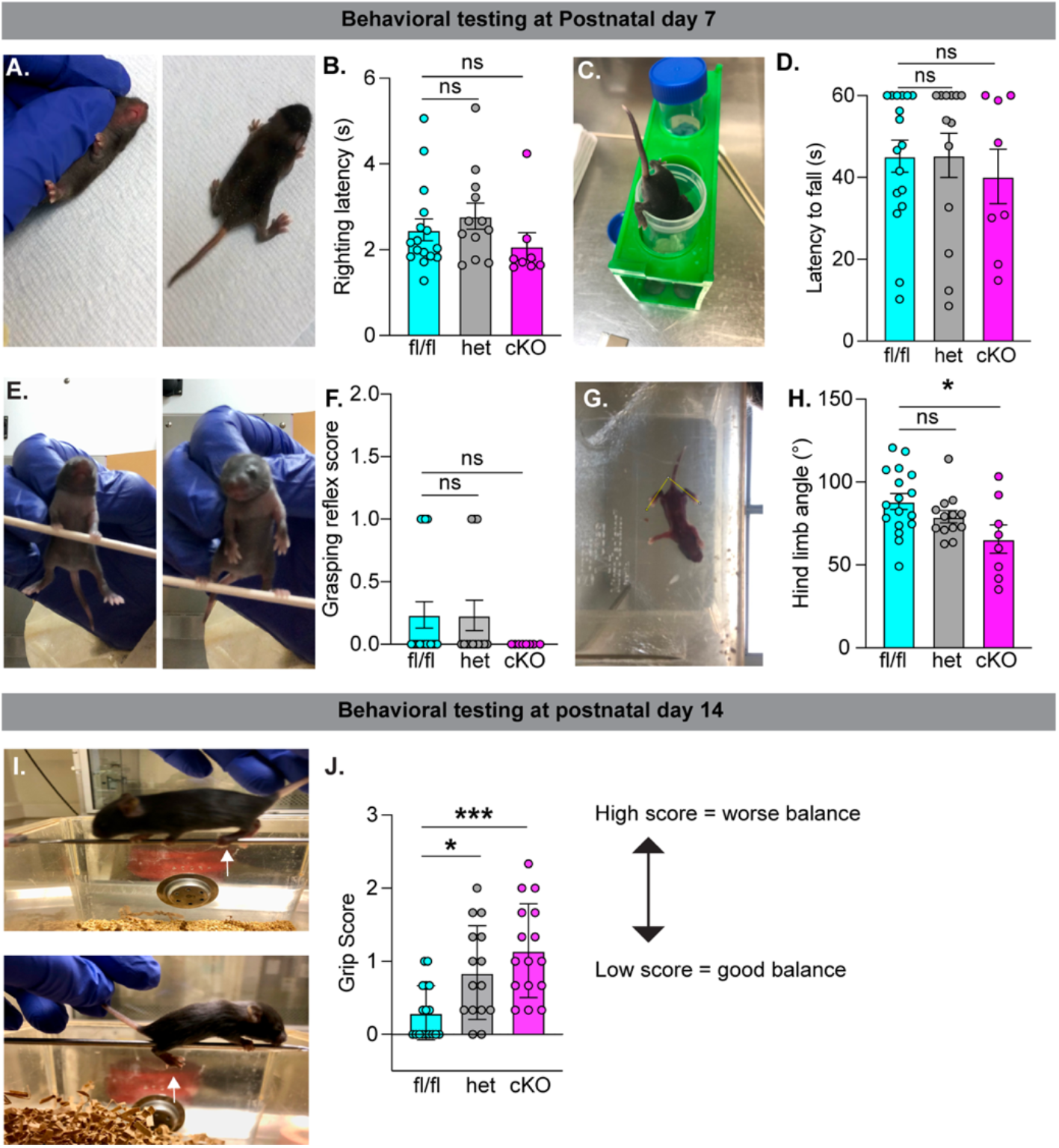
NaV1.6 is required in sensory neurons for motor behaviors in a developmentally dependent manner. (**A** to **H**) Behavioral testing on mice age P7, N=17 NaV1.6^fl/fl^, N= 13 NaV1.6^het^, and N= 8 NaV1.6^cKO^. (**A**)Representative images of righting reflex before (left) and after (right). (**B**) Latency for mice to right themselves was quantified. NaV1.6^het^ (grey, p=0.686), NaV1.6^cKO^ (magenta, p=0.655) compared to NaV1.6^fl/fl^ (cyan). (**C**) Representative image of hind limb strength assay. (**D**) Latency to fall was quantified. NaV1.6^het^ (p=0.999), NaV1.6^cKO^ (p=0.797) compared to NaV1.6^fl/fl^. (**E**) Representative images of grip reflex assay on forelimbs (left) and hindlimbs (right). (**F**) Mice were assayed on grip reflex. NaV1.6^het^ (p=0.999), NaV1.6^cKO^ (p=0.351) compared to NaV1.6^fl/fl^. (**G**) Representative image of hind limb angle quantification. (**H**) Quantification of mean hindlimb angle. NaV1.6^het^ (p=0.389), NaV1.6^cKO^ (p=0.021) compared to NaV1.6^fl/fl^. Behavioral testing on mice age P14. N=17 NaV1.6^fl/fl^, N=15 NaV1.6^het^, and N=16 NaV1.6^cKO^ (**I**) Representative images from limb coordination assay. Mice were scored based their ability to grasp the metal grate (top picture shows successful grasp, bottom picture shows foot slip). (**J**) Quantification of limb coordination score in P14 mice. N=17 NaV1.6^fl/fl^, N= 17 NaV1.6^het^, and N= 16 NaV1.6^cKO^. NaV1.6^het^ (p=0.022), NaV1.6^cKO^ (p=0.0002) compared to NaV1.6^fl/fl^. A one-way ANOVA with Tukey’s Post-hoc comparison was used to determine statistical significance.

**Fig. S3.**
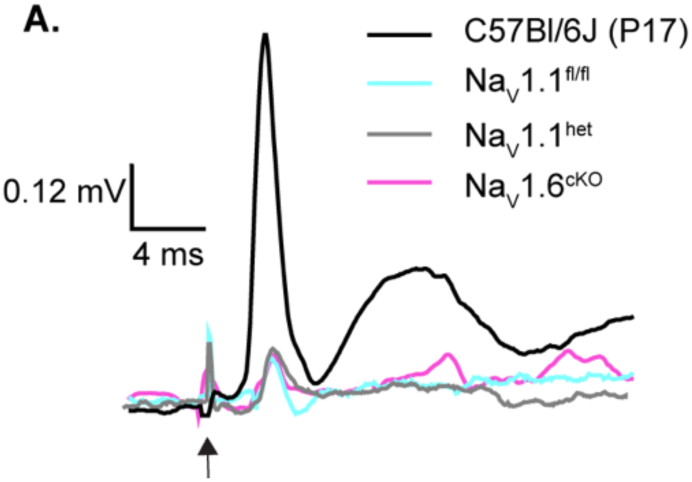
Recordings from Pirt^Cre^;NaV1.1 mice at late postnatal development. (**A**) P14 to P18 monosynaptic responses from NaV1.1^fl/fl^ (cyan), NaV1.1^het^ (grey), NaV1.1^cKO^ (magenta), and C57Bl/6J (black). Recordings from conditional mouse line exhibited small monosynaptic responses compared to age matched C57Bl/6J mice that were not reliably quantifiable.

**Fig. S4.**
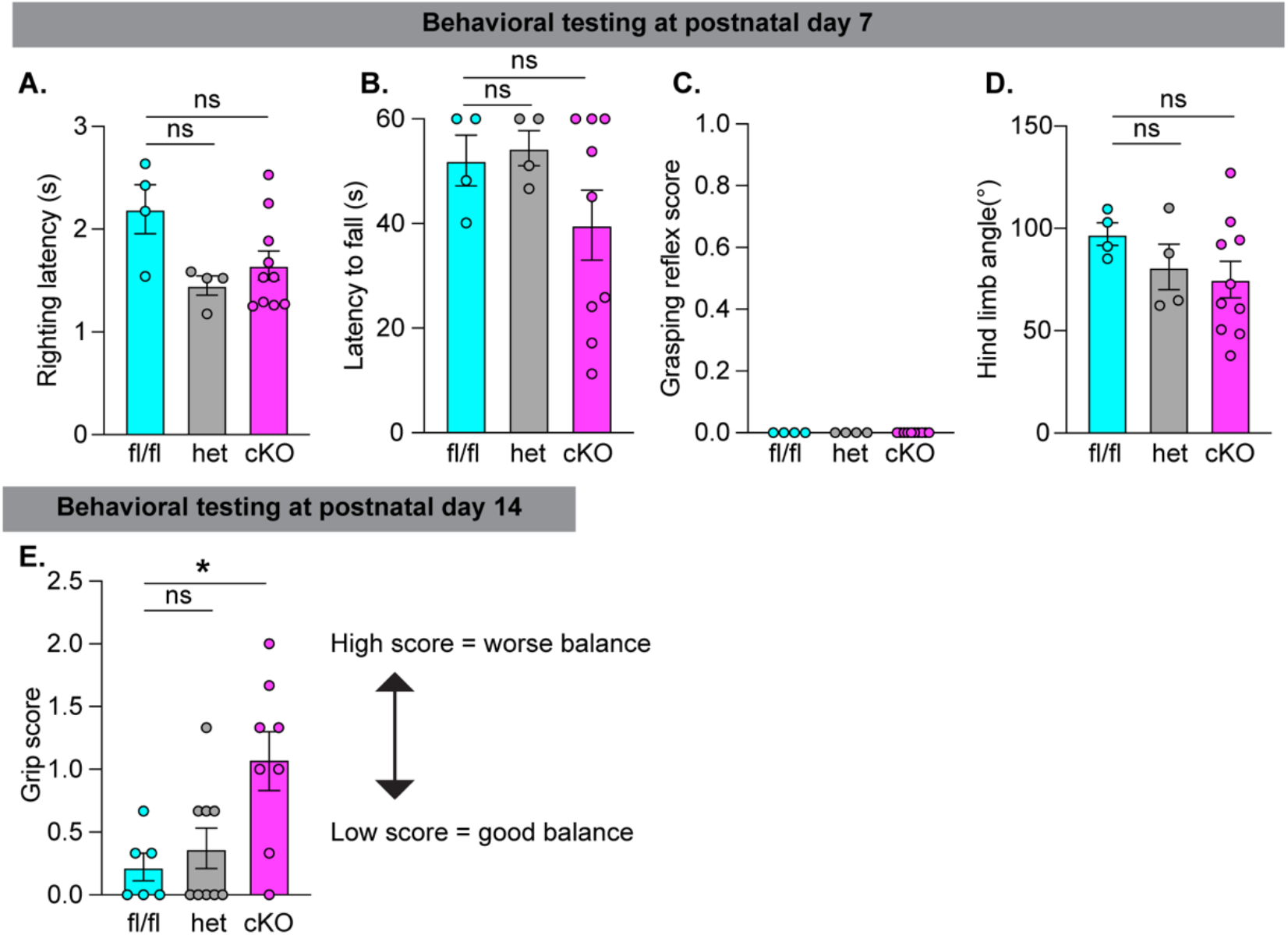
NaV1.1 in sensory neurons is required for motor function in late postnatal development. **(A)** to **D)** Behavioral testing on mice age P7. N=4 NaV1.1^fl/fl^, N= 4 NaV1.1^het^, and N= 9-10 NaV1.1^cKO^. (**A**) Latency for mice to right themselves was quantified. NaV1.1^het^ (grey, p=0.058), NaV1.1^cKO^ (magenta, p=0.142) compared to NaV1.1^fl/fl^ (cyan). (**B**) Latency to fall was quantified. NaV1.1^het^ (p>0.999), NaV1.1^cKO^ (p=0.715) compared to NaV1.1^fl/fl^. (**C**) Mice were assayed on grip reflex. All measured values between genotypes. (**D**) Quantification of mean hindlimb angle. NaV1.1^het^ (p=0.637), NaV1.1^cKO^ (p=0.906) compared to NaV1.1^fl/fl^. (**E**) Quantification of limb coordination score in P14 mice. N=6 NaV1.1^fl/fl^, N= 9 NaV1.1^het^, and N= 8 NaV1.1^cKO^. NaV1.1^het^ (p=0.845), NaV1.1^cKO^ (p=0.015) compared to NaV1.1^fl/fl^. A one-way ANOVA with Tukey’s Post-hoc comparison was used to determine statistical significance.

**Fig. S5.**
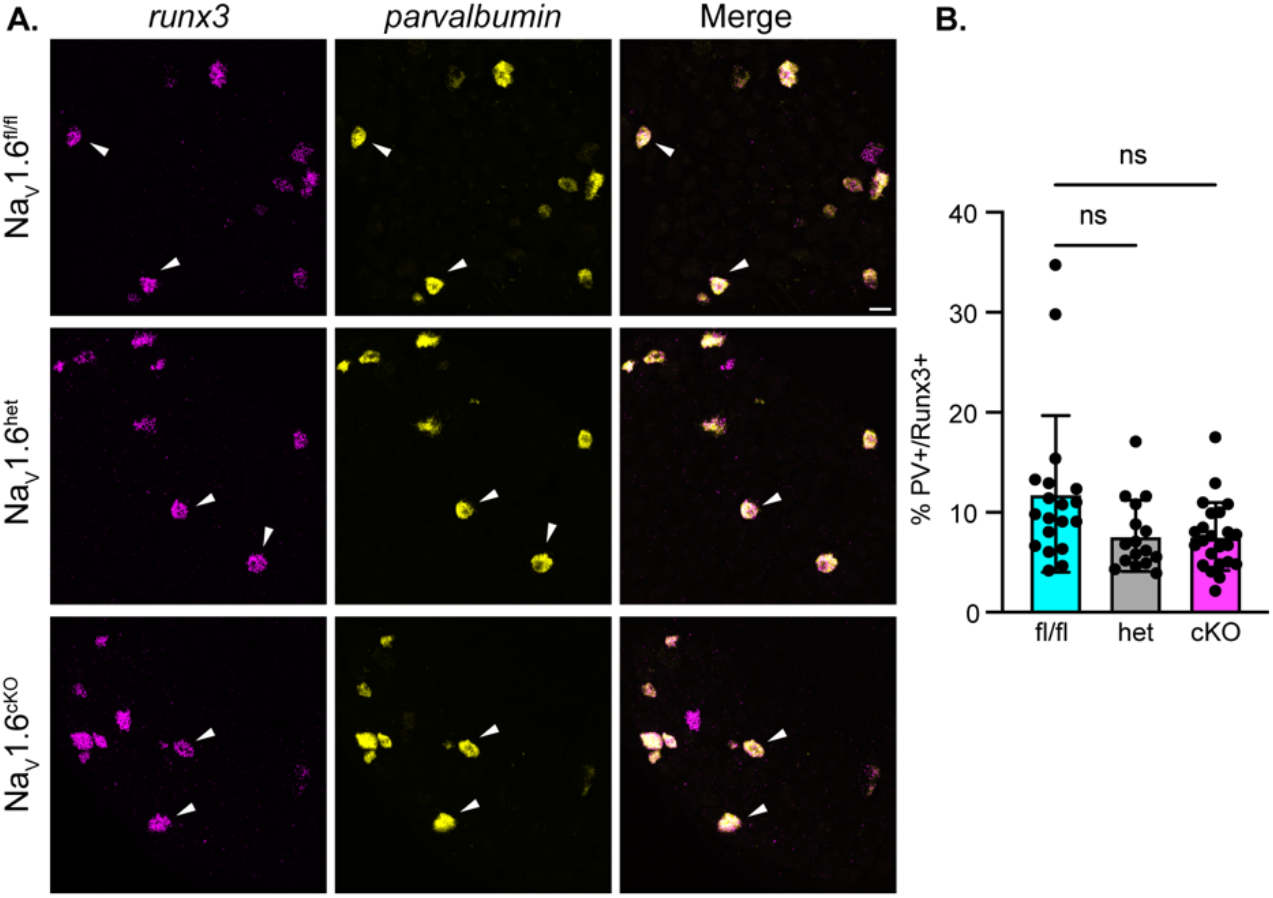
The number of proprioceptors in DRG is unaffected due to loss of NaV1.6 in sensory neurons. (**A**)Representative confocal images of NaV1.6^fl/fl^ (**top**), NaV1.6^het^ (**middle**), and NaV1.6^cKO^ (**bottom**) adult dorsal root ganglion (DRG) neuron section (25μm). Sections were hybridized with probes targeted against parvalbumin (Pvalb, yellow) and Runx3 (magenta). (**B**) Quantification of the percentage of Pvalb+/Runx3+ neurons per genotype. Each dot represents a single DRG section. Images were acquired with a 40x, 0.9 NA water immersion objective. A Kruskal-Wallis test with Dunn’s Post-hoc comparison was used to determine statistical significance. NaV1.6^het^ p=0.0694, NaV1.6^cKO^ p=0.0511 compared to NaV1.6^fl/fl^ . N=3 mice for each genotype. NaV1.6^fl/fl^ n = 19, NaV1.6^het^ n=16, NaV1.6^cKO^ n =22 sections.

**Fig. S6.**
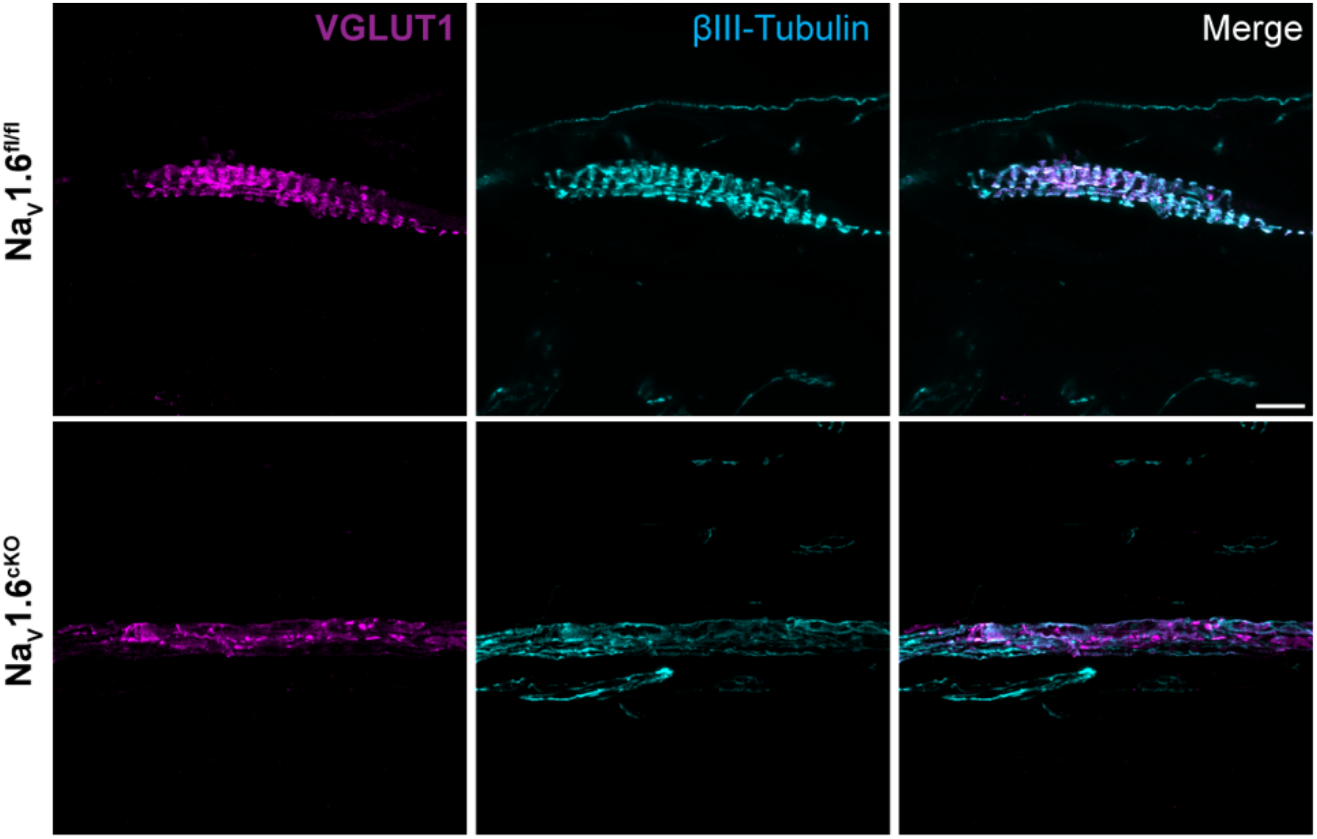
βIII-Tubulin and VGLUT1 labeling in muscle spindles are similar in mice NaV1.6^fl/fl^ and NaV1.6^cKO^ mice. Representative confocal images of muscle spindles from NaV1.6^fl/fl^ (**top**) and NaV1.6^cKO^(**bottom**) adult extensor digitorum longus muscle. Muscle spindle afferents are labeled were colabeled with VGLUT1 (magenta) and βIII-Tubulin (cyan) antibodies. Images were acquired with 60x, 1.4 NA oil immersion objective. Scale bar set to 25 μm.

**Fig. S7.**
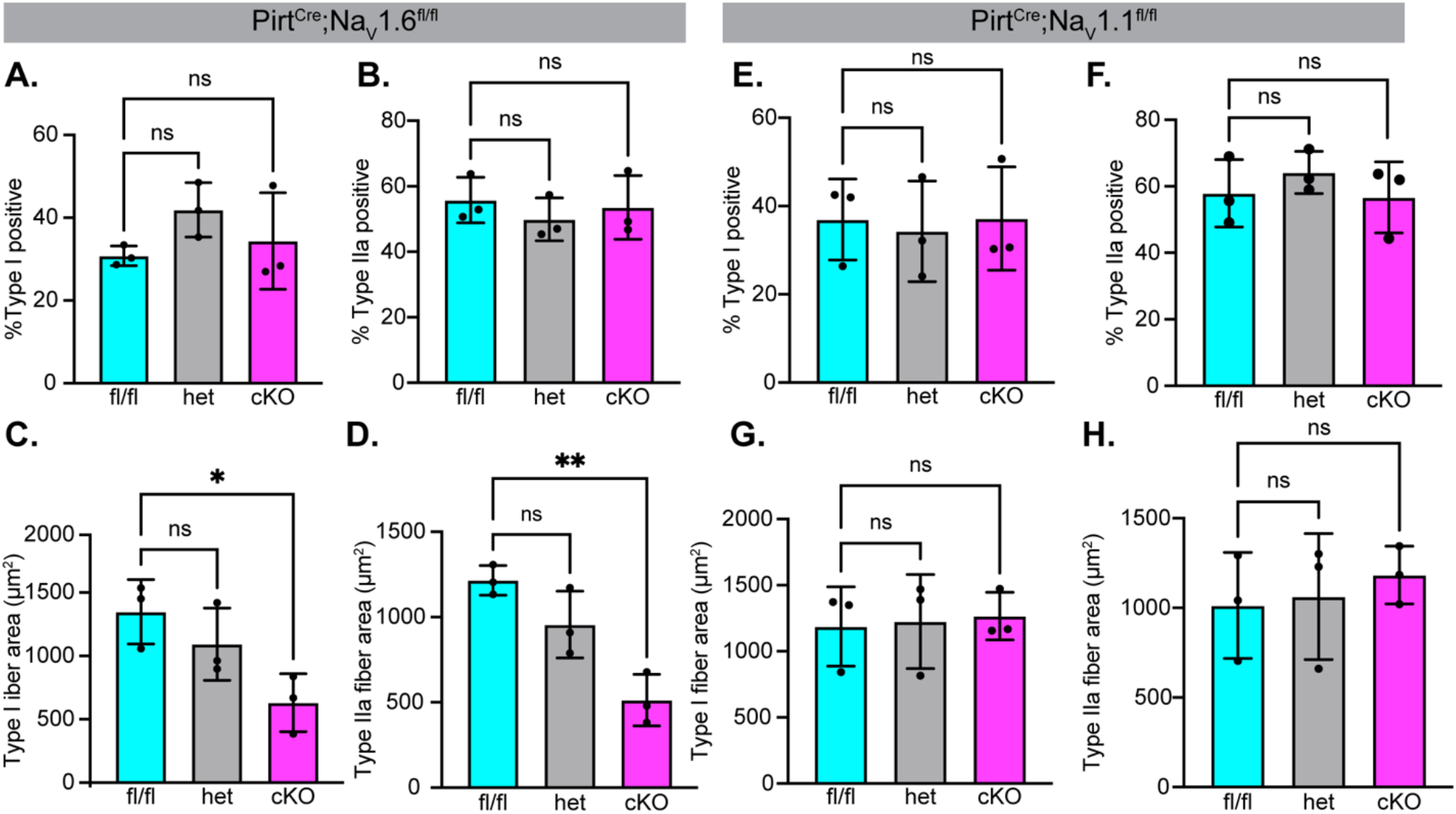
Muscle fiber type is unaffected due to loss of NaV1.6 or NaV1.1 in sensory neurons. (**A** to **D**) Quantification of average muscle fiber type in NaV1.6^fl/fl^ (cyan), NaV1.6^het^ (grey), and NaV1.6^cKO^ (magenta). (**A**) Percentage of type I positive muscle fibers, NaV1.6^het^ p=0.2217, NaV1.6^cKO^ p=0.8060 compared to NaV1.6^fl/fl^. (**B**) Type II positive muscle fibers, NaV1.6^het^ p=0.5861, NaV1.6^cKO^ p=0.9170 compared to NaV1.6^fl/fl^. (**C**) Type I fiber area, NaV1.6^het^ p=0.4209, NaV1.6^cKO^ p=0.0250 compared to NaV1.6^fl/fl^. (**D**) Type II fiber area, NaV1.6^het^ p=0.1376, NaV1.6^cKO^ p=0.0023 compared to NaV1.6^fl/fl^. (**E** to **H**) Quantification of average muscle fiber type in NaV1.1^fl/fl^ (cyan), NaV1.1^het^ (grey), and NaV1.1^cKO^ (magenta). (**E**) Percentage of type I positive muscle fibers, NaV1.1^het^ p=0.9358, NaV1.1^cKO^ p=0.9995 compared to NaV1.1^fl/fl^. (**F**) Type II positive muscle fibers, NaV1.1^het^ p=0.9662, NaV1.1^cKO^ p=0.6959 compared to NaV1.1^fl/fl^. (**G**) Type I fiber area, NaV1.1^het^ p=0.9819, NaV1.1^cKO^ p=0.9236 compared to NaV1.1^fl/fl^. (**H**) Type II fiber area, NaV1.1^het^ p=0.6435, NaV1.1^cKO^ p=0.9799 compared to NaV1.1^fl/fl^. A one-way ANOVA with Dunn’s post hoc comparison was used to determine statistical significance. Each dot represents a single animal. N=3 for each genotype.

**Fig. S8.**
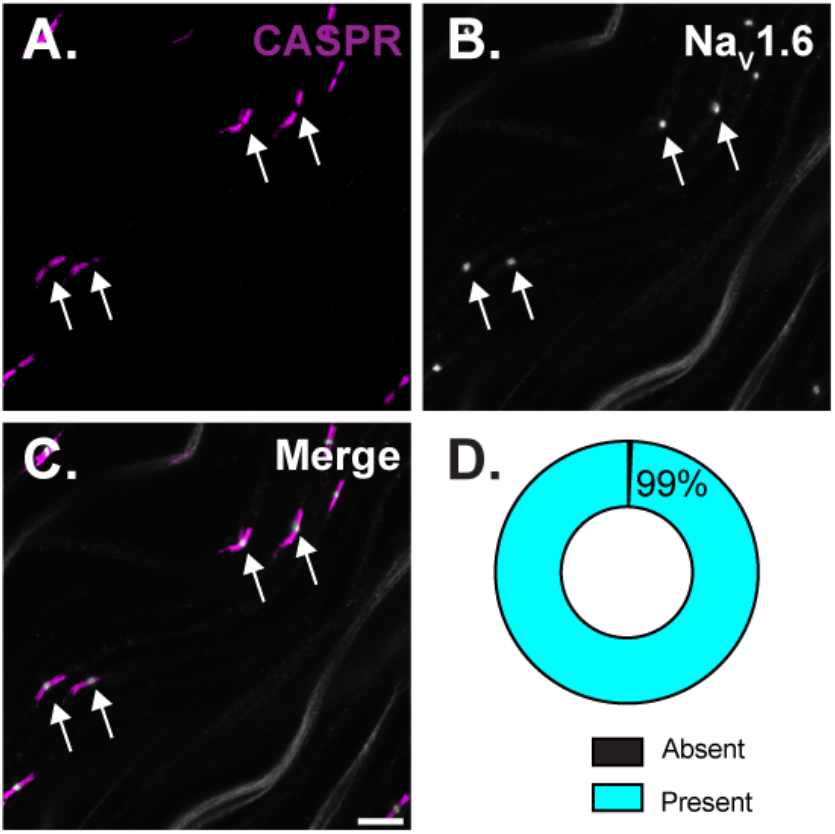
NaV1.6 highly expressed at nodes of Ranvier sensory axons. (**A** to **C**) Representative images for sensory nodes of Ranvier were identified via CASPR (**A**) and NaV1.6 (**B**) immunoreactivity. (**D**) Quantification of the percentage of nodes of Ranvier that express NaV1.6. n=247 nodes, N=3 mice. Scalebar=10μm

**Fig. S9.**
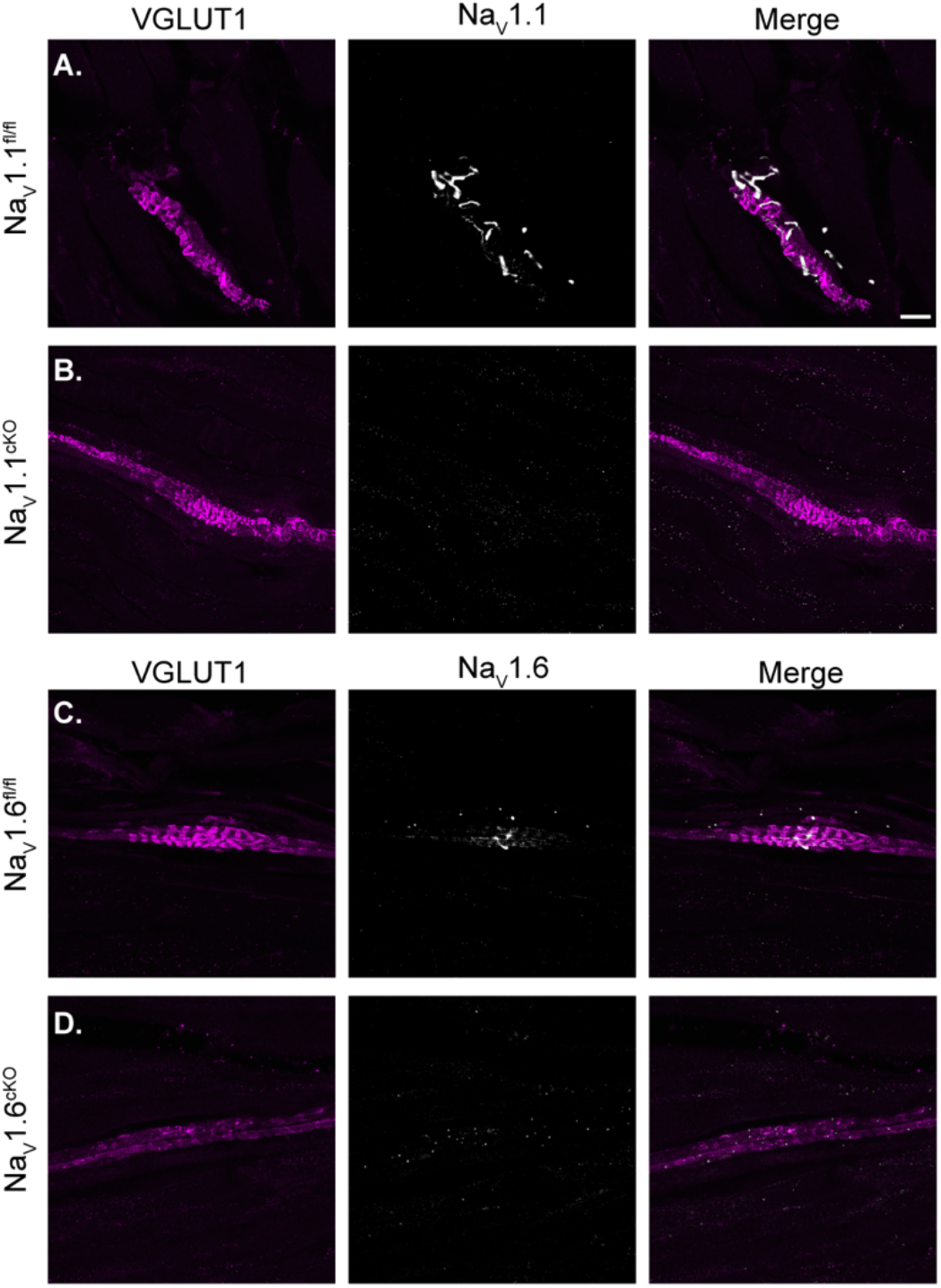
Validation of NaV antibodies targeting NaV1.1 and NaV1.6. Representative confocal images of muscle spindles from NaV1.1^fl/fl^ (**A**), NaV1.1^cKO^(**B**), NaV1.6^fl/fl^ (**C**) and NaV1.6^cKO^ (**D**) adult extensor digitorum longus muscle. VGLUT1 (magenta) labels muscle spindle sensory endings. Grey scale represents corresponding NaV channel isoform. Scale bar set to 20 μm. Images were acquired with a 60x, 1.4 NA oil immersion objective.

**Fig. S10.**
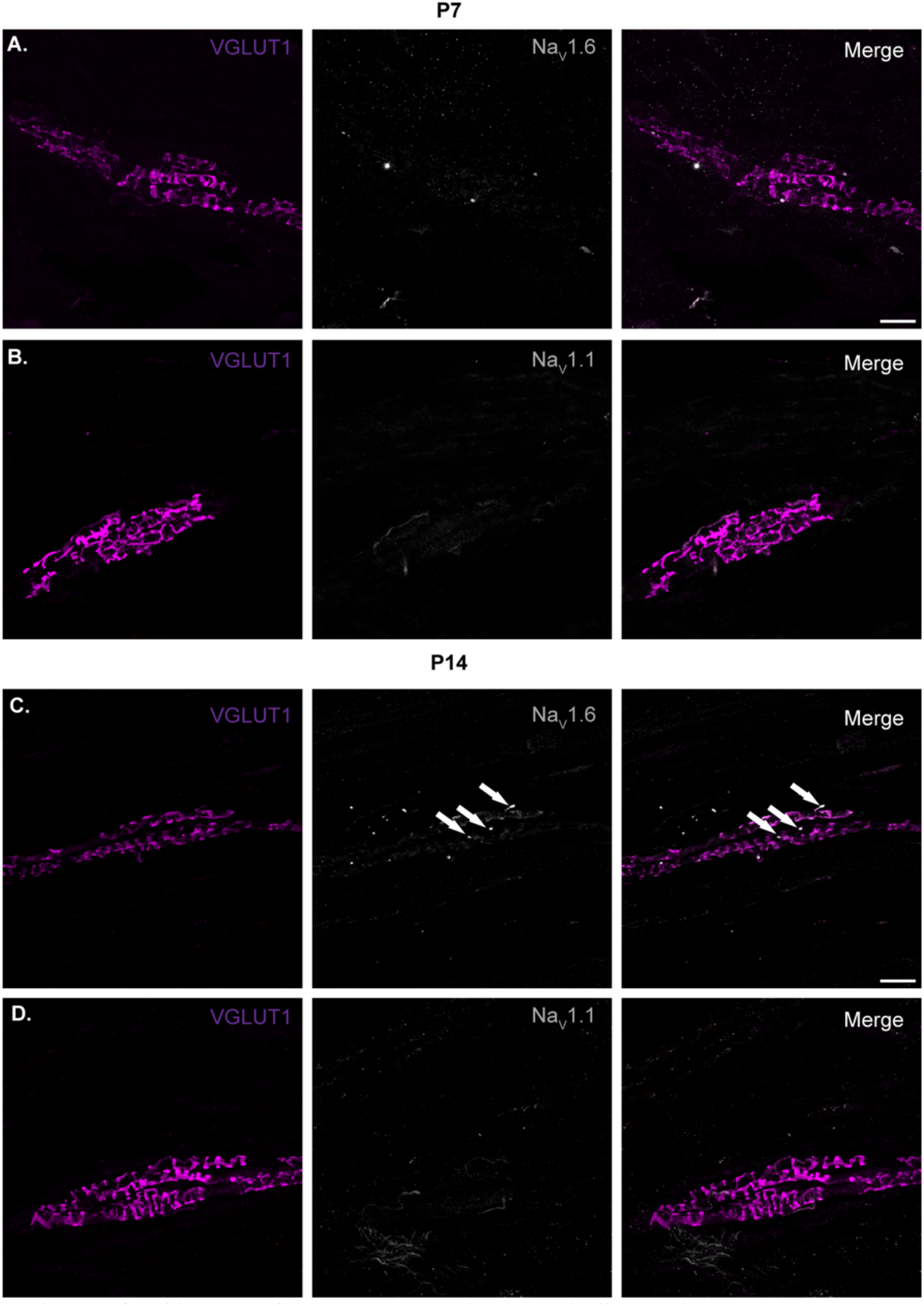
NaV labeling in muscle spindles throughout postnatal development. Representative confocal images of NaV1.6 (**A**, 3= mice, 4= spindles) and NaV1.1 (**B**, 4= mice, 6= spindles) labeling in muscle spindles from extensor digitorum longus muscle from mice at postnatal day 7. Images of NaV1.6 (**C**, 3= mice, 7= spindles, arrows denote clusters of NaV1.6) and NaV1.1 (**D**, 7= mice, 16= spindles) in mice at postnatal day 14. Images were acquired with a 60x, 1.4 NA oil immersion objective. All tissue was collected from C57BL/6J mice. Scalebar=25 μm.

## Supplementary Tables

**Table S1:** Within-genotype analysis of the monosynaptic reflex response in the NaV1.6 mouse line across postnatal development. P-values obtained from within genotype, across development, statistical analyses (Two-way ANOVA) of the monosynaptic reflex response in NaV1.6fl/fl, NaV1.6het, and NaV1.6cKO mice.

|  | P6-P11 vs. P14-P18 |  |  |
| --- | --- | --- | --- |
|  | Nav1.6<sup>fl/fl</sup> | Nav1.6<sup>het</sup> | Nav1.6<sup>cKO</sup> |
| Latency | 0.019** | 0.0515 | 0.8039 |
| Threshold | 0.3536 | 0.2799 | 0.0027** |
| Monosynaptic peak amplitude | 0.6501 | <0.0001**** | 0.0027** |
| Polysynaptic peak amplitude | 0.1764 | 0.0002*** | 0.0032** |
| FWHM | 0.1848 | 0.8767 | 0.0002*** |

## Supplementary Movies

**Movie S1.**
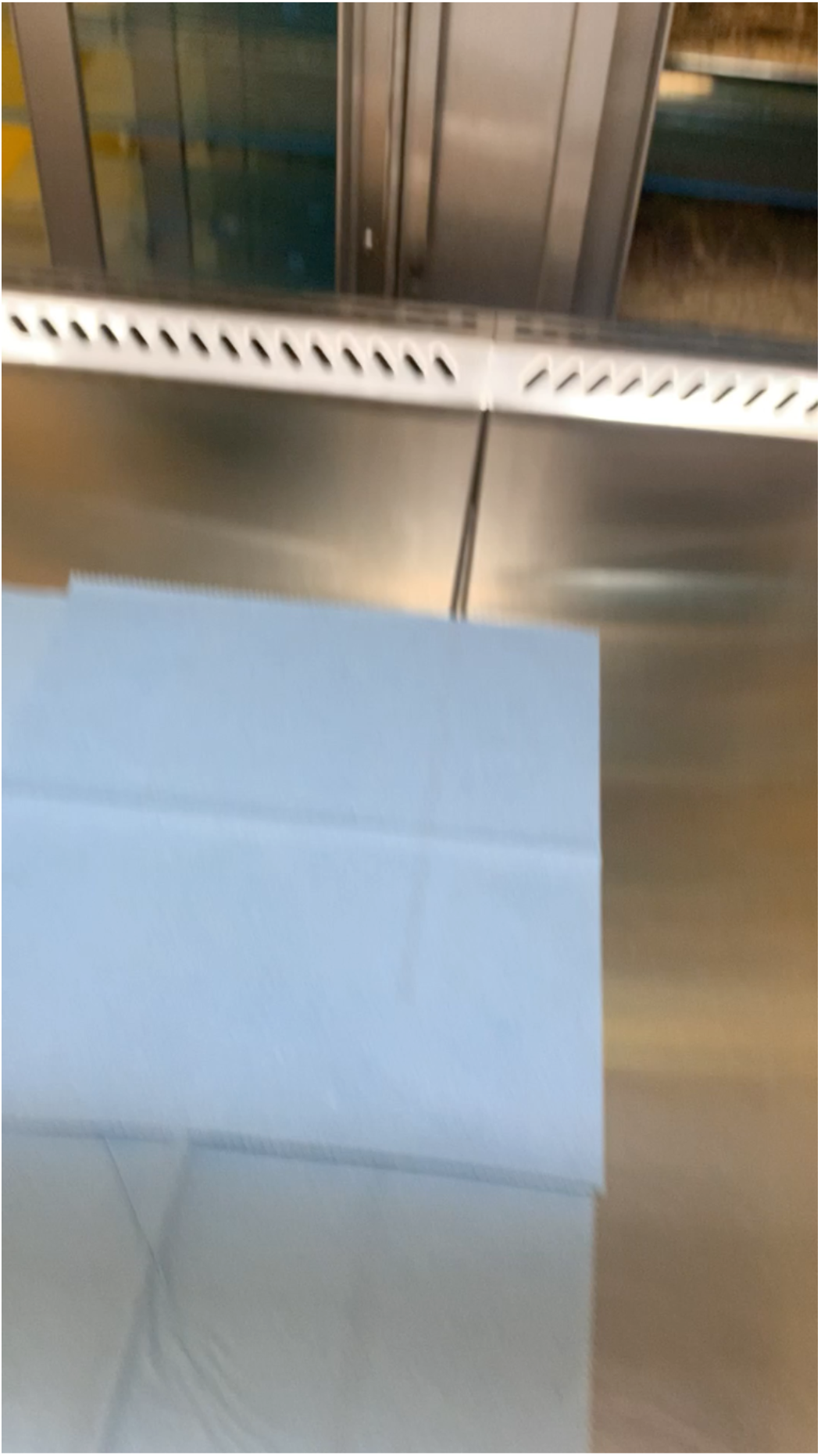
Complete loss of limb coordination in NaV1.6cKO mice when suspended by tails.

**Movie S2.**
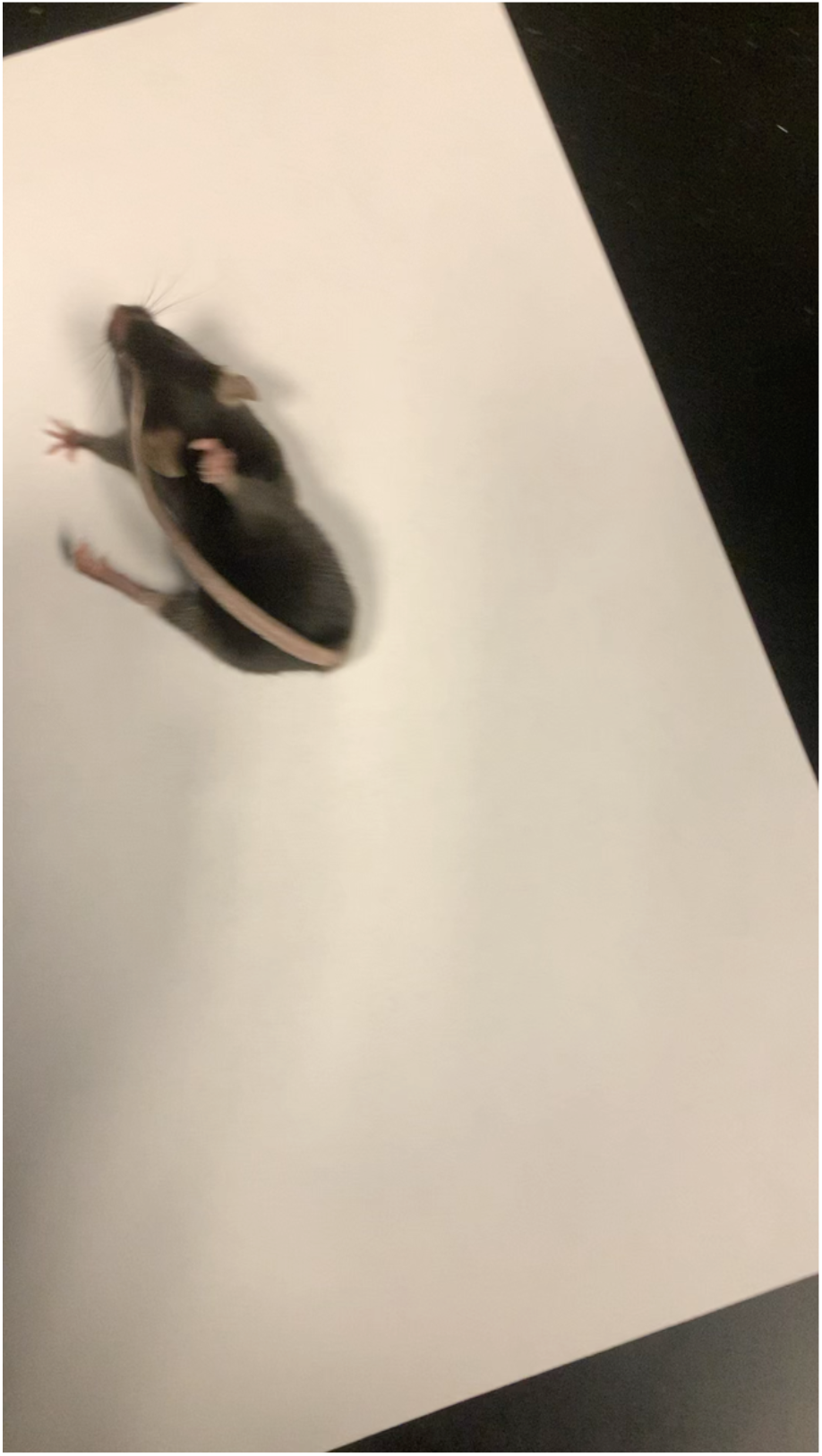
An example of a NaV1.6cKO mouse unable hindlimbs or tail to for normal walking behaviors.

